# A reductase in the lipid metabolism at cross roads between cuticular wax production and jasmonic acid-mediated defenses in maize

**DOI:** 10.1101/2022.08.10.503514

**Authors:** Jiong Liu, Lu Li, Christelle AM Robert, Baozhu Li, Shan He, Zhilong Xiong, Wenjie Chen, Jiasheng Bi, Guanqing Zhai, Siyi Guo, Hui Zhang, Jieping Li, Shutang Zhou, Xi Zhang, Chun-Peng Song

## Abstract

Cuticular wax is the first physical layer to protect plants from herbivore infestations. Although wax and jasmonic acid (JA) biosynthesis are the two most studied branches of lipid metabolism, the mechanism how cuticular wax production modulates plant chemical defenses is unclear. Here, we show that the maize (*Zea mays*) *GL8* gene, encoding 3-ketoacyl reductase during fatty acid elongation in the biosynthesis of very long chain fatty acids (VLCFA), functions as a turning point between wax production and JA-mediated defenses. The fall armyworm (*Spodoptera frugiperda*) larvae consumed more tissue but gained a lower performance on *gl8/Ye478* mutant plants. *gl8/Ye478* mutant displayed higher JA-mediated defenses constitutively, and also more inducible by herbivore stimulation. The comprehensive transcriptomic and lipidomic analyses further demonstrated that Zm*GL8* mutation up-regulates the JA biosynthesis pathway by promoting the accumulation of lysophospholipids and modulation of galactolipid synthase genes *ZmDGD* and *ZmMGD*. The phenotypic and transcription comparisons of the maize and Arabidopsis wax-deficient mutants suggest a conserved wide-spread trade-off between wax production and chemical defense in both 18:3 and 16:3 plants. These results illustrate a critical role of VLCFA metabolism as a switch to control the balance between cuticular wax physical barrier and JA-mediated chemical defenses during plant-herbivore coevolution history.

## Introduction

In the natural environment, plants are usually attacked by a various of herbivore insects, which seriously threaten plant growth and crop production (Agrawal, 2011; Fernandez et al., 2021). The hydrophobic extracellular biopolymers of cuticular wax was evolved by land plants about 450 million years ago, which is an important adaptation to survive under the arms-races between plants and herbivores (Eigenbrode and Espelie, 1995; Wang et al., 2015; Kong et al., 2020). Through affecting herbivore attachment, feeding and oviposition, cuticular wax is at multiple levels as the first physical layer to protect plant aboveground tissues from herbivore attacks (Eigenbrode and Jetter, 2002; Lewandowska et al., 2020).

Cuticular wax is an important group of plant lipids, including a complex mixture of very long-chain fatty acids (VLCFA) and their derivatives (Jetter et al., 2008). VLCFA starts with a *de novo* production of fatty acyl-acyl carrier proteins (ACPs) of C_16_ and C_18_ structure in plastids (Samuels et al., 2008). ACP and free fatty acids are released by acyl-ACP thioesterase (FATA and FATB), and esterified to acyl-Coenzyme A (CoA) by long-chain acyl-CoA synthetase (LACS) (Li-Beisson et al., 2013; Lee and Suh, 2015; Hegebarth et al., 2017). Acyl-CoAs are the very beginning point for both VLCFA formation and the phytohormone jasmonic acid (JA) biosynthesis (Joubes et al., 2008; Hurlock et al., 2014; Lewandowska et al., 2020; Wan et al., 2020).

JAs, a group of oxylipins from oxygenated fatty acid products, are critical for plants to successfully thwarting herbivore attacks (Borrego and Kolomiets, 2016). Oxylipins are generated from C_18_ structure unsaturated fatty acids, such as linoleic or linolenic acid via lipoxygenase (LOX) (Borrego and Kolomiets, 2016; Hu et al., 2021). Linoleic and linolenic acids are derived from galactolipids monogalactosyl diacylglycerol (MGDG) and digalactosyl diacylglycerol (DGDG) of chloroplast membranes (Yu et al., 2020). MGDG and DGDG are formed from hydrolysis of diacylgycerol and phospholipids by phospholipases (Mongrand et al., 1998; Okazaki and Saito, 2014). Based on the galactolipid metabolism, plants can be defined as 18:3 and 16:3 plants. Maize (*Zea mays* L.), wheat (*Triticum aestivum* L.) and rice (*Oryza sativa* L.) are typical 18:3 plants, and *Arabidopsis thaliana* and spinach (*Spinacia oleracea*) are 16:3 plants, which use 18:3 or 16:3 structure lipids as start material for MGDG and DGDG biosynthesis (Kelly et al., 2016). Even there is a potential inner connection between cuticular wax and JA biosynthesis in plant lipid metabolism, it is unknown whether and how cuticular wax constrains plant JA-manipulated chemical defenses against herbivores on the genetic level.

Maize is one of the most important crops in the world, and its cuticular wax is an important agronomic trait. Based on the advanced researches on cuticular wax of Arabidopsis (Li et al., 2013), 12 maize mutants have been reported involved in cuticular wax biosynthesis, transport, mediation, and tolerance to stresses. In particular, *ZmGL6*, *ZmSRL5*, *ZmFDL1, ZmGL1*, *ZmGL15*, *ZmFDH1* and *ZmFAE1* can manipulate maize drought tolerance (Yu et al., 2017; Li et al., 2019; Castorina et al., 2020; Pan et al., 2020), and only *ZmGL11* was shown to be involved in resistance to biotic stress, the powdery mildew fungus (*Blumeria graminis* f.sp. *hordei*) (Hansjakob et al., 2011). Whether wax biosynthesis genes modulate maize resistance to herbivory has not been studied in detail so far.

Here, based on the water-loss phenotype, we identified a ‘*wal1*’ mutant line from Ye478 background maize EMS-mutagenized library, whose water-loss and cuticular wax deficiency phenotypes are caused by *ZmGL8* mutation. Our study demonstrated that *ZmGL8* can balance maize physical and chemical defenses against the fall armyworm caterpillars. By combining transcriptomic and lipidomic analyses, we highlighted potential interactions between *ZmGL8,* maize lipid metabolism, and herbivore resistance. Using a JA-biosynthesis inhibitor treatment, we confirmed that *ZmGL8* regulated chemical defenses is JA-dependent. Finally, we show that wax deficiency also enhances chemical defense levels in Arabidopsis, suggesting a possible widespread phenomenon in both 18:3 and 16:3 plants. Collectively, these experiments unravel a novel function of cuticular wax on herbivore chemical defenses and a trade-off between physical and chemical defenses mediated by a single gene locus.

## Results

### Cuticular wax deficiency promotes maize chemical defense against herbivores

From the 2,000 maize EMS-mutagenized lines in Ye478 background, we found a line with clear water-loss phenotype, which was named as ‘*wal1*’ mutant. The *wal1* mutant had a higher water loss rate, brighter green leaf, the ability to hold water drops and a quasi-absence of wax crystals on its leaf surface under scanning by electron microscope (SEM) (Figure 1A;Supplementary Figure S1A). The cuticular wax analysis revealed reduced amounts of very long chain fatty acids, aldehydes, and alkanes in *wal1* mutant (Supplementary Figure S1B). However, no difference in leaf cell inner structure was observed between WT and *wal1*mutant maize plants through a transmission electron microscope (TEM) (Supplementary Figure S1C). Map-based cloning and bulked segregate analysis-sequencing method adjusted from the protocol described by (Wang et al., 2019) showed the targeted gene of the *wal1*mutant was *ZmGL8* (Zm00001d017111), which changed the 200^th^ basic group G into A in the first exon of the fifth chromosome (Supplementary Figure S2). This result was confirmed by allele validation of *gl8* mutant from B73 background EMS Mutagenesis library and the segregation statistics of the F1 generation (Supplemental Table S1). *ZmGL8* is known to encode 3-ketoacyl reductase, which is the essential part of elongases of very long chain fatty acids (Xu et al., 2002). Then, we named the *wal1* mutant to *gl8/Ye478* for subsequent experiments.

**Figure 1.**
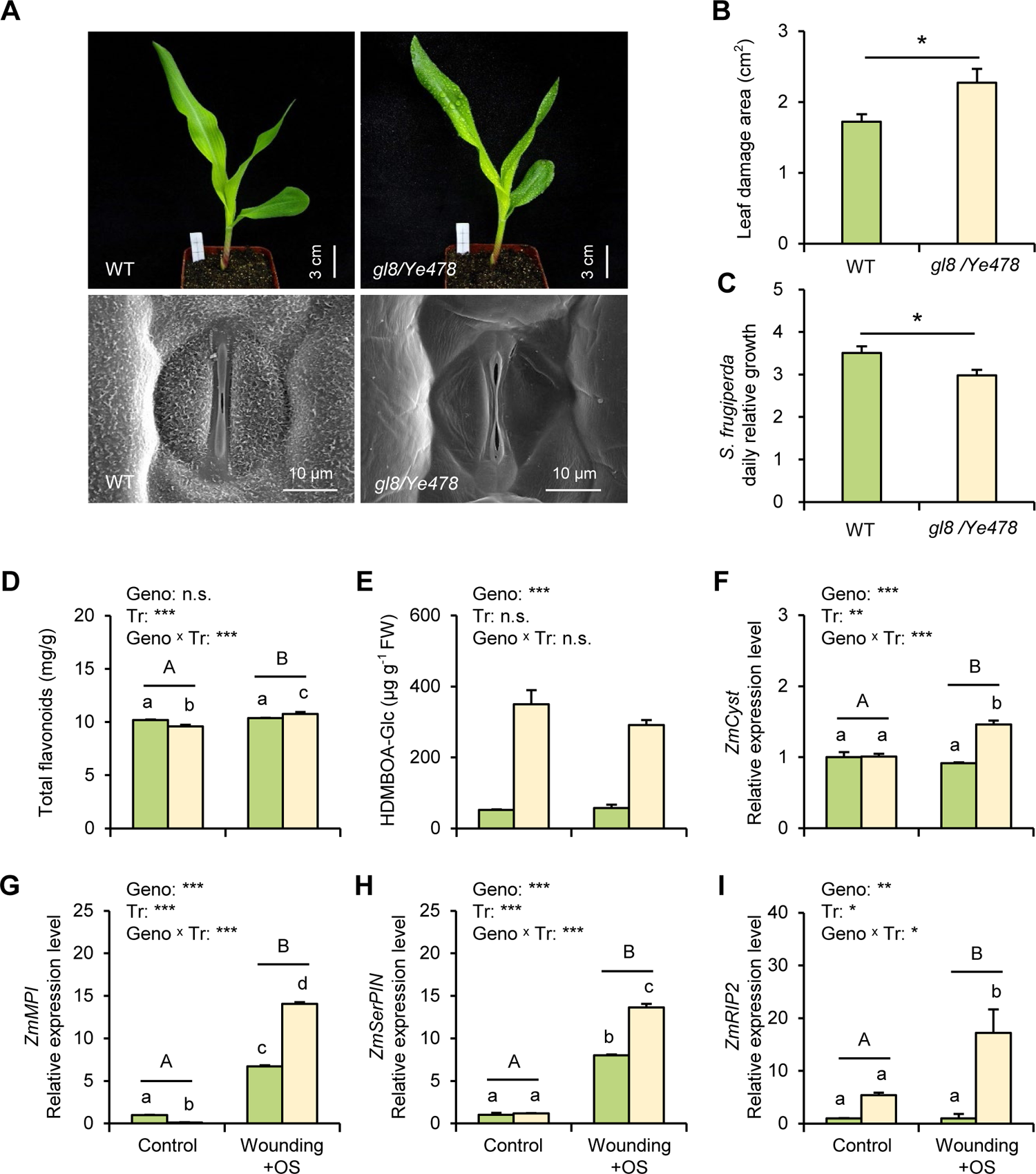
*gl8/Ye478* mutant is more resistant to herbivores. **(A)** Cuticular wax phenotype of *gl8/Ye478* mutant and wild-type (WT) plants after water spraying on leaf surface (Bar = 3 cm) and observations under SEM (×6000 magnification, Bar = 10 μm). **(B)** Leaf damage (Mean ± SE, n = 14-15) of *Spodoptera frugiperda* caterpillars feeding on *gl8/Ye478* mutant and WT maize plant leaves for 8 hours. **(C)** Daily relative growth (Mean ± SE, n = 20) of *S. frugiperda* caterpillars feeding on *gl8/Ye478* mutant and WT maize plants for 72 hours. **(D)** to **(E)** Concentrations of total flavonoids (Mean ± SE, n = 3) and HDMBOA-Glc (Mean ± SE, n = 4) of *gl8/Ye478* mutant and WT maize plants under control and wounding + oral secretion (OS) induction treatments. **(F)** to **(I)** Gene expression (Mean ± SE, n = 3) of the putative proteinase inhibitors *ZmMPI*, *ZmSerPIN*, and *ZmCyst*, and the insecticidal ribosome-inactivating protein *ZmRIP2*. Green bars: WT plants; Yellow bars: *gl8/Ye478* mutant plants. Asterisks indicate significant differences between genotypes, treatments or the two factor interactions (*\*P* < 0.05, *\*\*P* < 0.01, *\*\*\*P* < 0.001). Lower case letters indicate the significant differences of pairwise comparisons, and uppercase letters indicate significant differences between control and wounding + OS treatments.

Interestingly, we found the fall armyworm *Spodoptera frugiperda* larvae consumed larger amounts of *gl8/Ye478* mutant leaves (Figure 1B), but grew slower than on WT maize plants (Figure 1C). To test whether *gl8/Ye478* mutant plants have stronger chemical defense against *S. frugiperda* larvae, we analyzed the concentrations of two major maize secondary metabolites and the expression of four defense-related genes. We used artificial wounding and *S. frugiperda* larva oral secretion (OS) to simulate herbivory to avoid the possible herbivore consumption bias of WT and *gl8/Ye478* mutant plants. The wax deficient mutant *gl8/Ye478* had overall similar levels of flavonoids to WT plants, but wounding + OS induced a stronger response in *gl8/Ye478* mutant (Figure 1D). The *gl8/Ye478* mutant constitutively produced higher concentrations of the benzoxazinoid HDMBOA-Glc than WT plants, although wounding + OS did not induce the compound in any of the two genotypes (Figure 1E). *gl8/Ye478* mutant plants had higher expression levels of the putative proteinase inhibitors *ZmMPI*, *ZmSerPIN, ZmCyst*, and insecticidal ribosome-inactivating protein *ZmRIP2* than WT plants under wounding + OS stimulations (Figure 1F-I). Besides, total soluble sugar and protein concentrations were similar in both genotypes (Supplementary Figure S3).

### *ZmGL8* up-regulated jasmonic acid biosynthesis and related herbivore defense pathways

To have an overall view of how wax biosynthesis deficiency influence maize chemical defense against herbivores, we conducted transcriptome analysis of WT and *gl8/Ye478* mutant plants under wounding + OS and control conditions. A principal component analysis (PCA) showed that the herbivore simulation drastically influenced the transcriptional level of maize plants (PC1, 63%), and the difference between WT and *gl8/Ye478* mutant plants was also obviously (PC2, 16%) (Figure 2A). The DEGs between WT and *gl8/Ye478* mutant plants were 1524 (with 774 up-regulated and 750 down-regulated) and 1738 (with 1125 up-regulated and 613 down-regulated) under control and herbivore-induced treatments, respectively (Supplementary Figure S4). The Kyoto Encyclopedia of Genes and Genomes (KEGG) enrichment analysis showed that these DEGs were mostly involved in herbivore-defense related pathways and most of them are related to or interact with JA-biosynthesis pathway, such as the phenylpropanoid biosynthesis, phenylalanine metabolism, linoleic acid and alpha-linolenic acid metabolism pathways (Howe and Schilmiller, 2002; Barros and Dixon, 2020) (Figure 2B). Marker genes involved in JA biosynthesis, PAL family genes and benzoxazinoid biosynthesis were mostly upregulated in *gl8/Ye478* mutant plants (Figure 2C). This pattern was further confirmed by qRT-PCR for selected markers (Supplementary Figure S5). Thus, *ZmGL8* mutation probably promotes maize chemical defenses by up-regulating JA biosynthesis and related downstream secondary metabolite metabolism.

**Figure 2.**
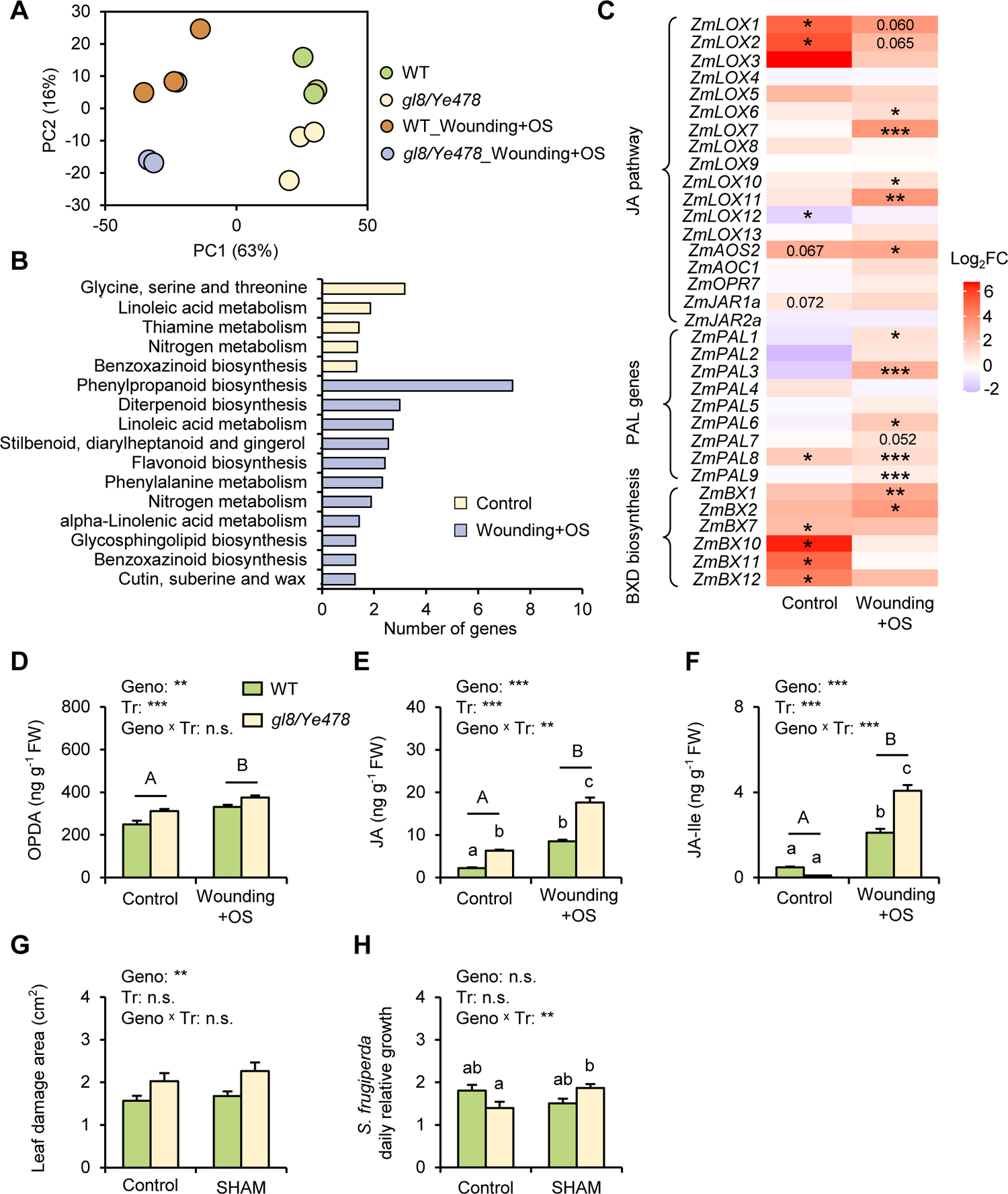
*gl8/Ye478* increased resistance to herbivory is mediated by the JA-pathway in maize. **(A)** to **(C)** Transcriptome analysis of *gl8/Ye478* mutant and WT maize plants under control and wounding + OS induction treatments (n = 3). **(A)** Principal component analysis (PCA) of all the gene expression levels from transcriptome analysis; **(B)** Kyoto Encyclopedia of Genes and Genomes (KEGG) enrichment analysis for pathways enriched in *gl8/Ye478* mutant; **(C)** Heat map of JA pathway, PAL family, and benzoxazinoid biosynthesis related genes in *gl8/Ye478* mutant compared with WT plants (n = 3). The color gradient represents the relative sequence abundance. Numbers in the color key indicate log_2_ fold change (FC). **(D)** to **(F)** JA and its derivative concentrations of *gl8/Ye478* mutant and WT maize plants under control and wounding + OS induction treatments. **(D)** OPDA, **(E)** JA and **(F)** JA-Ile concentrations in related treatments (Mean ± SE, n = 4). **(G)** Leaf damage (Mean ± SE, n = 13-15) by *S. frugiperda* caterpillars feeding for 8 hours on JA-biosynthesis inhibitor salicylhydroximic acid (SHAM)-treated *gl8/Ye478* mutant and WT maize plant leaves. **(H)** Daily relative growth (Mean ± SE, n = 18-21) of *S. frugiperda* caterpillars feeding on SHAM treated maize plants for 72 hours. Asterisks indicate significant differences between genotypes, treatments or the two factor interactions (*\*\*P* < 0.01, *\*\*\*P* < 0.001). Lower case letters indicate the significant differences of pairwise comparisons, and uppercase letters indicate significant differences between control and wounding + OS treatments.

### JAs play a key role in *ZmGL8* up-regulated chemical defenses in maize

To clarify the role of JAs in *gl8/Ye478* mutant up-regulated defenses, we analyzed the accumulation of compounds from the JA-biosynthesis pathway. The concentrations of 12-oxo-phytodienoic acid (OPDA) and JA were constitutively higher in *gl8/Ye478* mutant plants (Figure 2D-F). Similarly, abscisic acid (ABA) and salicylic acid (SA), but not 1-aminocyclopropane-l-carboxylic acid (ACC, a biosynthetic precursor of ethylene) concentrations were more elevated in *gl8/Ye478* mutant plants than in WT, but SA was with a similar concentration in both genotypes (Supplementary Figure S6). Wounding + OS triggered a higher induction of OPDA, JA, jasmonyl-_L_-isoleucine (JA-Ile) and ABA in *gl8/Ye478* than in WT (Figure 2D-F; Supplementary Figure S6).

We then used the JA biosynthesis inhibitor salicylhydroximic acid (SHAM), to test the importance of JA biosynthesis on *ZmGL8*-mediated chemical defense differences. SHAM reduced the expression of the marker genes involved in JA biosynthesis (*ZmLOX10* and *ZmAOS*), benzoxazinoid biosynthesis (*ZmBX1* and *ZmBX10/11*), terpenoid biosynthesis (*ZmIGL*, *ZmTPS2, ZmTPS3,* and *ZmTPS10*), as well as in proteinase inhibitor production (*ZmMPI* and *ZmRIP2*) in *gl8/Ye478* mutant plants (Supplementary Figure S7). Surprisingly, SHAM induced the expression of *ZmLOX10*, *ZmLOX11*, *ZmAOS*, *ZmMPI, ZmTPS2*, *ZmTPS3*, and *ZmTPS10,* and only reduced the expression of *ZmIGL* in WT plants (Supplementary Figure S7). After treating maize plants by SHAM, *S. frugiperda* larvae damaged area was not influenced, as the feeding surface was still larger in *gl8/Ye478* mutant than in WT (Figure 2G). However, SHAM addition resulted in a higher growth rate of *S. frugiperda* larvae on *gl8/Ye478* mutant than on non-treated plants, even though the growth of the caterpillars was not influenced by SHAM in WT plants (Figure 2H).

### *ZmGL8* modulates the lipid metabolism

As both plant cuticular wax and JA biosynthesis are important branches of the lipid metabolism, we conducted a lipidomic analysis for WT and *gl8/Ye478* mutant plants under control and wounding + OS conditions. The PCA result showed a significant difference between WT and *gl8/Ye478* mutant plants (PC1, 42%), but only a slight difference was induced by herbivore stimulation (Figure 3A). There were 186 lipids significantly influenced by wounding + OS, in which 112 lipids were involved in both WT and *gl8/Ye478* mutant, 50 lipids were specific to WT and 24 lipids to *gl8/Ye478* mutant (Supplementary Figure S8). Consistently with the transcriptome analysis, the lipid metabolism KEGG enrichment analysis also showed that linoleic acid and alpha-linolenic acid metabolism pathways were significantly induced in *gl8/Ye478* mutant. Additionally, the glycerolipid and glycerophospholipid metabolisms were also significantly influenced by the *ZmGL8* mutation (Figure 3B).

**Figure 3.**
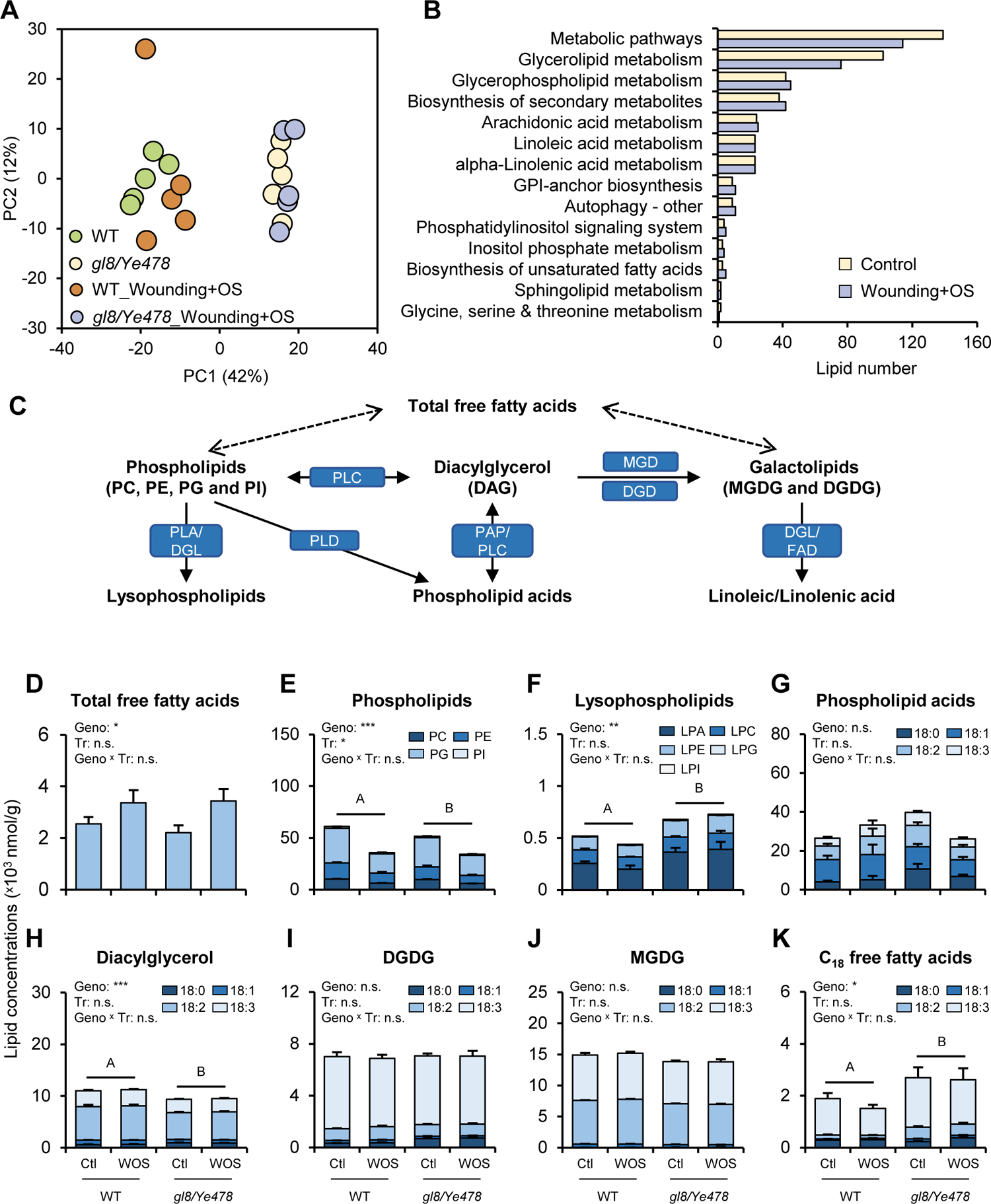
*ZmGL8* modulates the maize lipid metabolism. **(A)** to **(B)** Lipidomic analysis of *gl8/Ye478* mutant and WT maize plants under control and wounding + OS induction treatments (n = 5). **(A)** Principal component analysis (PCA) of all detected lipid concentrations; **(B)** Kyoto Encyclopedia of Genes and Genomes (KEGG) enrichment analysis for pathways enriched in *gl8/Ye478* mutant. **(C)** Lipid metabolism pathways related to JA pathway. The enzymes are shown in dark blue squares. **(D)** to **(K)** Lipid concentrations of **(D)** total free fatty acids, **(E)** phospholipids, **(F)** lysophospholipid, **(G)** Phospholipid acids, **(H)** diacylglycerol, **(I)** digalactosyl diacylglycerol (DGDG), **(J)** monogalactosyl diacylglycerol (MGDG) and **(K)** C_18_ free fatty acid (FFA) in *gl8/Ye478* mutant and WT maize plants under wounding + OS induction. PC: phosphatidylcholine, PE: phosphatidylethanolamine, PG: phosphatidylglycerol, PI: phosphatidylinositol, LPA: lysophosphatidic acid, LPC: lysophosphatidyl choline, LPE: lysophosphatidyl ethanolamine, LPG: lysophosphatidyl glycerol, LPI: lysophosphatidyl inositol, Ctl: control, WOS: Wounding + OS, 18:0-18:3: lipids with this structure. Asterisks indicate significant differences between genotypes, treatments or the two factor interactions (*\*P <* 0.05, *\*\*P <* 0.01, *\*\*\*P <* 0.001). Uppercase letters indicate significant differences between *gl8/Ye478* mutant and WT maize plants.

Then, we analyzed the lipids related to JA biosynthesis in detail (Figure 3C). Although the wax contents were dramatically reduced (Supplementary Figure S1B), the total free fatty acid concentrations were increased in the *gl8/Ye478* mutant than WT plants (Figure 3D). Free fatty acids are related with phospholipid and galacotolipid transformation pathways (Okazaki and Saito, 2014). In phospholipids related pathway, concentrations of phosphatidylcholine (PC), phosphatidylethanolamine (PE), phosphatidylglycerol (PG) and phosphatidylinositol (PI) were significantly reduced in *gl8/Ye478* mutant plants (Figure 3E), while that of lysophosphatidic acid (LPA), lysophosphatidyl choline (LPC), lysophosphatidyl ethanolamine (LPE), lysophosphatidyl glycerol (LPG), lysophosphatidyl inositol (LPI) and phospholipid acid (PA) with 18:0 structure were much induced in *gl8/Ye478* mutant plants (Figure 3F-G). Transcriptome analysis showed phospholipase genes, including *ZmPLD3*, *ZmPLD7*, *ZmPLD10* and *ZmDGL5*, were significantly induced in *gl8/Ye478* mutant. *ZmPLA2* also up-regulated more in *gl8/Ye478* mutant under herbivore-induced conditions (Supplementary Figure S8, B and C). In galactolipid related pathway, the concentrations of diacylglycerol (DAG), monogalactosyl diacylglycerol (MGDG) and digalactosyl diacylglycerol (DGDG) with 18:0 structure were higher in *gl8/Ye478* mutant, while MGDG and DGDG concentrations with 18:1 structure were higher in WT plants (Figure 3, H-J). DGDG synthase *ZmDGD* was down-regulated, but MGDG synthases *ZmMGD* and *ZmDGL5* were up-regulated (Supplementary Figure S8, B and C). Finally, C_18_ free fatty acids, such as oleic acid, linoleic acid and linolenic acid were significantly or with a trend to accumulate more in *gl8/Ye478* mutant than WT plants (Figure 3K). To investigate whether the high saturated free fatty acids can be converted into low saturated linoleic acid and linolenic acid by desaturase, we analyzed the expression levels of fatty acid desaturase FAD family and found the *ZmFAD2* and *ZmFAD6* were up-regulated in *gl8/Ye478* mutant plants (Supplementary Figure S8, B and C). All the details for statistics were summarized in Supplemental Table S3.

### Wax-mediated defense modulation is conserved in maize and Arabidopsis

To investigate the conservation of wax biosynthesis deficiency up-regulated chemical defense, we collected maize and Arabidopsis mutants with genes in different positions of wax biosynthesis pathway (Figure 5). We first observed the wax deficiency phenotype of the chosen mutants. Similar to *gl8/Ye478* mutant, *gl1/B73*, *gl6/B73*, *gl8/B73* and *gl14/B73* mutants also have more bright green leaves and can hold water on their leaf surface. The SEM observation of the leaf surface showed there was no wax crystal on *gl1/B73* and *gl8/B73* mutants, while *gl6/B73* and *gl14/B73* mutants had few wax crystals but still much less than WT plants (Supplementary Figure S9). However, because of the trichomes on Arabidopsis leaf surface, we could only observe the wax deficiency by SEM. The phenotype in Arabidopsis was not as obvious as in maize, but still had fewer wax crystals on mutants than Col-0 (Supplementary Figure S10).

To check the chemical defense levels of different maize and Arabidopsis wax deficiency mutants, we tested the *S. frugiperda* performance, defense-related secondary metabolite production and gene expression levels. Consistent with the results found in *gl8/Ye478* mutant, *S. frugiperda* larvae fed more on the maize wax deficient mutants *gl1/B73*, *gl6/B73* and *gl8/B73* (Figure 4A), but there was no difference of *S. frugiperda* larva growth rate among mutants and WT maize plants (Figure 4B). Flavonoid concentrations were constitutively lower in *gl6/B73, gl8/B73,* and *gl14/B73* mutants, but wounding + OS triggered a stronger flavonoid induction in *gl1/B73* and *gl8/B73* mutants than in WT (Figure 4C). Interestingly, wounding + OS did not induce flavonoids in *gl6/B73* and *gl14/B73* mutants (Figure 4C). HDMBOA-Glc concentrations were higher in the mutants than in the WT plants, except in *gl8/B73* under wounding + OS condition, and Wounding + OS slightly decreased the concentrations of the benzoxazinoid in all genotypes (Figure 4D). Gene expression levels of JA-biosynthesis pathway marker genes, *ZmPLD* family genes, and *ZmMGD* were increased in mutant maize plants (Supplementary Figure S11).

**Figure 4.**
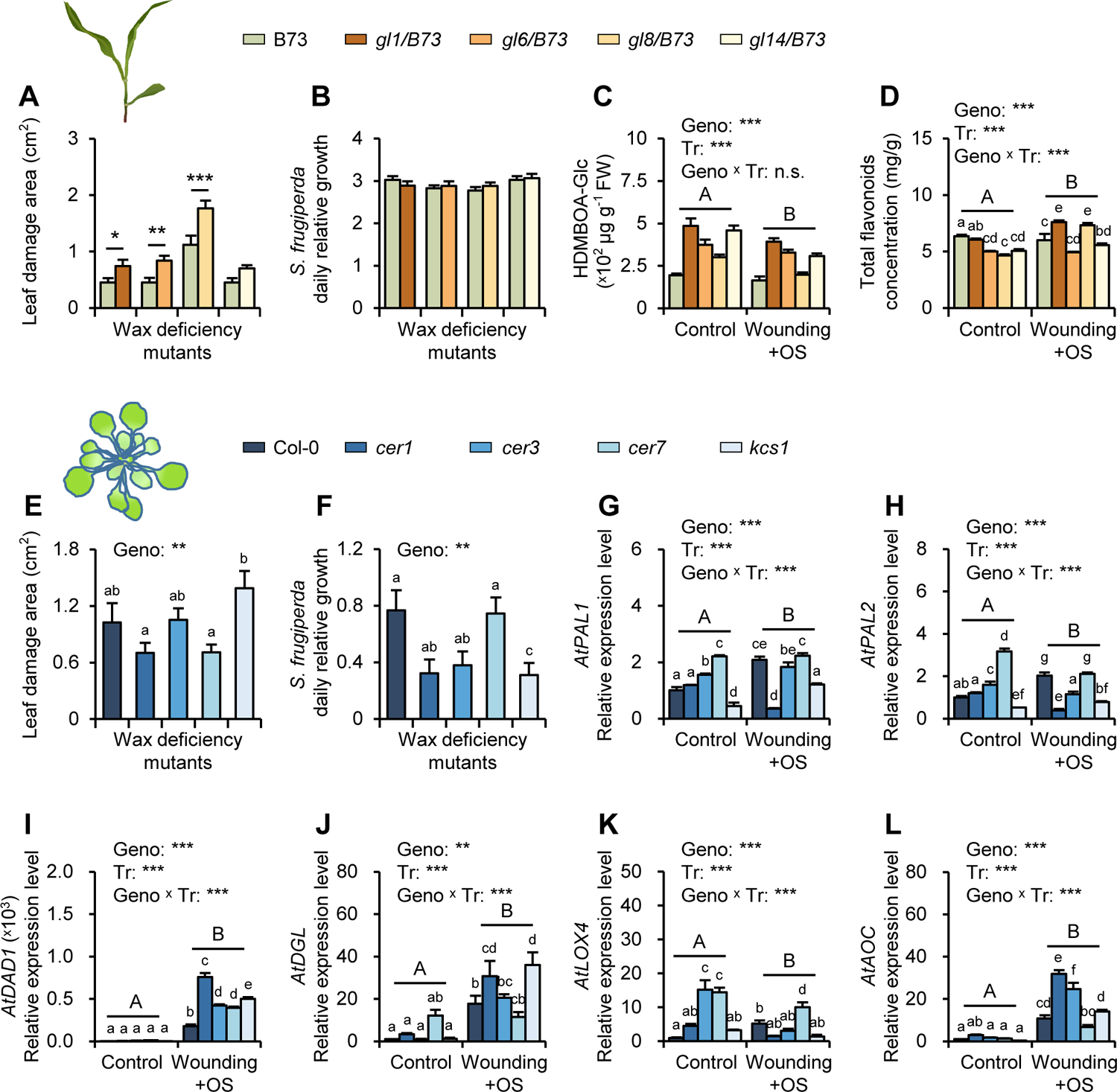
The JA-mediated increase in herbivore resistance is common in maize and Arabidopsis wax deficient mutants. **(A)** Leaf damage (Mean ± SE, n = 15) of *S. frugiperda* caterpillars feeding on different cuticular wax deficient mutants and WT maize plants for 8 hours. **(B)** Daily relative growth (Mean ± SE, n = 20) of *S. frugiperda* caterpillars feeding on different cuticular wax deficient mutants and WT maize plants for 72 hours. **(C)** to **(D)** Concentrations of total flavonoids (Mean ± SE, n = 3) and HDMBOA-Glc (Mean ± SE, n = 4) among different wax deficient mutants and WT maize plants under control and wounding + OS induction treatments. **(E)** Leaf damage (Mean ± SE, n = 14-15) of *S. frugiperda* caterpillars feeding on different cuticular wax deficient mutants and WT *A. thaliana* for 8 hours. **(F)** Daily relative growth (Mean ± SE, n = 8-10) of *S. frugiperda* caterpillars feeding on different cuticular wax deficienty mutants and WT *A. thaliana* plants for 72 hours. **(G)** to **(L)** Gene expression levels (Mean ± SE, n = 3) of JA pathway related genes in control and wounding + OS treatments of *A. thaliana*. Asterisks indicate significant differences between genotypes, treatments or the two factor interactions (*\*P* < 0.05, *\*\*P* < 0.01, *\*\*\*P* < 0.001). Lower case letters indicate the significant differences of pairwise comparisons, and uppercase letters indicate significant differences between control and wounding + OS treatments.

In Arabidopsis, a mutation in wax production (*cer1*, *cer3, cer6*, and *kcs1*) resulted in mutation-specific differences in leaf consumption and herbivore growth. In particular, *Atkcs1* mutant, whose mutation is also in the fatty acid elongation cycle as *Zmgl8* mutant (Lewandowska et al., 2020), yielded a slight, albeit not significant increase in leaf consumption (Figure 4E), and a decrease in herbivore growth (Figure 4F). A trend for a lower herbivore performance was further observed on *cer1*, *cer3*, but not *cer8* mutants (Figure 4F). Marker genes involved in the JA-biosynthesis pathway were mostly upregulated in wax deficient Arabidopsis lines (Figure 4 G-L). A mutation in the *AtCER* genes led to higher constitutive expression of *AtPAL1, AtPAL2* and *AtLOX4* (Figure 4G-H and K). Wounding + OS triggered a higher induction of *AtDAD1* for all mutants, *AtDGL* for *cer1* and *kcs1* mutants, *AtLOX4* for *cer8* mutants and *AtAOC* expressions in the *cer1*, *cer3* and *kcs1* mutants than in Col-0 (Figure 4I-L). In total, maize wax deficiency mutants showed stronger chemical defense than WT plants, while Arabidopsis wax deficiency could also up-regulate chemical defense but not as strong as maize plants.

## Discussion

Plants have evolved complex physical and chemical defense traits to cope with herbivore infestation. Here, we reveal that *ZmGL8* modulates a trade-off between wax production and JA-depended chemical defenses (Figure 5). The cuticular wax benefits the herbivore which, despite a reduced feeding, performs better on wax-producing leaves. Below, we discuss the potential mechanisms and the conservation of cuticular wax trade-off functions on plant chemical and physical defenses.

**Figure 5.**
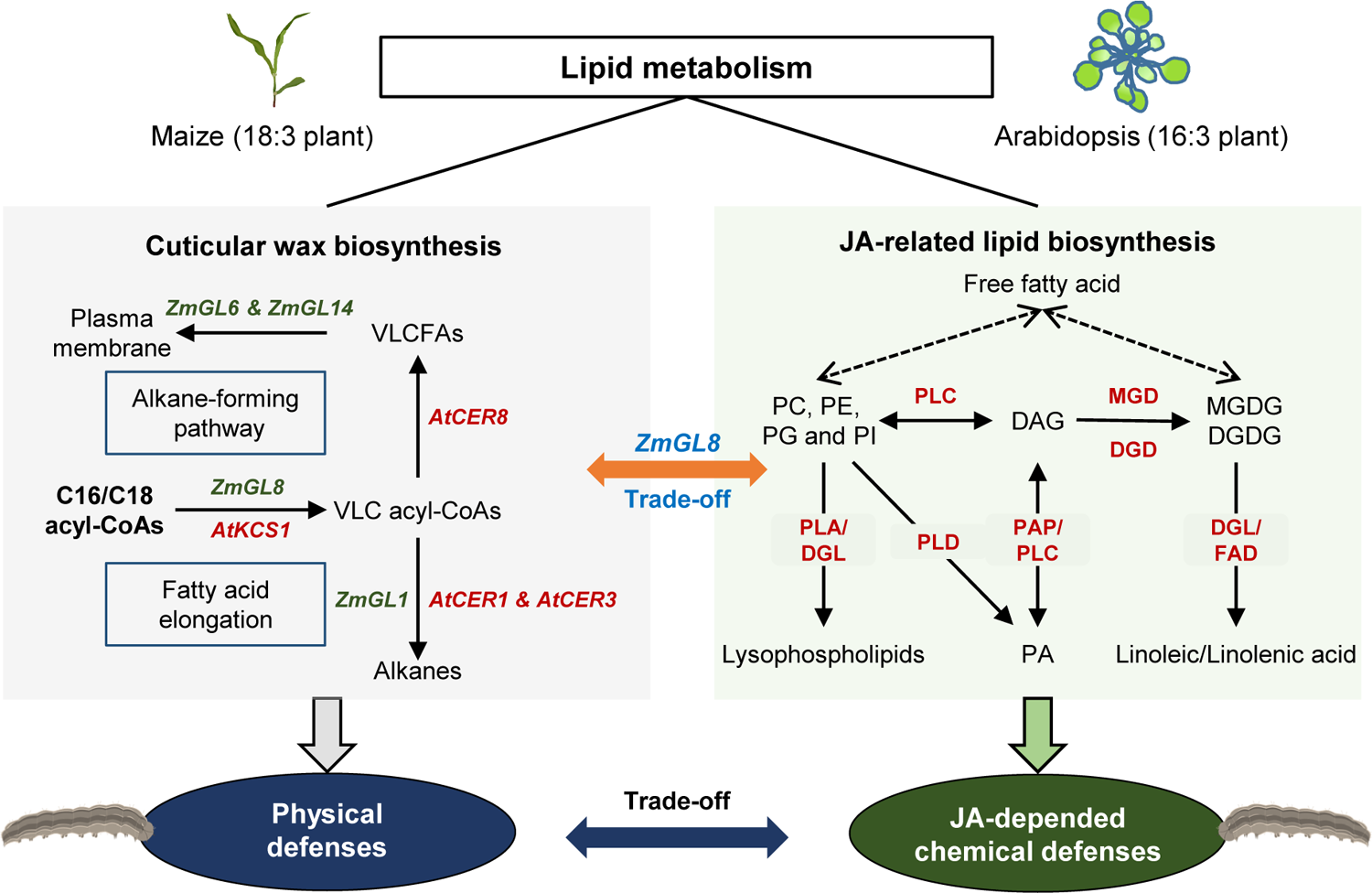
A model for potential mechanism of wax biosynthesis mediated physical and JA-depended chemical defenses against herbivores. The cuticular wax biosynthesis is composed by fatty acid elongation, alkane-forming pathway and very long chain fatty acid transformation to the plasma membrane, in which we selected *ZmGL8*, *ZmGL1*, *ZmGL6* and *ZmGL14* mutation maize plants and *AtKCS1*, *AtCER1*, *AtCER3* and *AtCER8* mutation Arabidopsis plants to test the conversation of wax deficiency mediated JA-related resistance in 16:3 and 18:3 plants. *ZmGL8* influence the pool of free fatty acids and subsequently remodeling plant lipid metabolism. For detail, *ZmGL8* mutation raises the transformation of phospholipids to lysophospholipids, free fatty acids and phospholipids, that are second messengers to promote JA pathway and downstream defenses; *ZmGL8* mutation can also increase JA biosynthesis through galactolipids derived linoleic and linolenic acid pathway. Thus, wax biosynthesis can mediate plant physical and chemical defenses trade-offs of plants under herbivore attacks.

Our results show that cuticular wax reduce herbivore damage, but increase herbivore performance. The damage reduction is consistent with the fact that wax represents a physical barrier for insect feeding. For example, removing wax mechanically from *Brassica* spp. can greatly increase beetle feeding, and younger leaves of *Eucalyptus* spp. are also protected by more complex wax microstructure than glossier mature leaves (Bodnaryk, 1992; Eigenbrode and Espelie, 1995). However, despite such a reduction in feeding, the herbivore grew better in presence of wax on the leaves, which may indicate a better fitness for herbivores (Veyrat et al., 2016). These observations together with ours may reflect either a better nutritive value as food for an herbivore or by lower defenses in wax-producing plants. While the first hypothesis can, at this stage, not be excluded, our data reported no differences in sugar and protein concentrations between WT and wax-deficient mutant maize plants. However, wax-producing plants exhibited lower constitutive JA-related defenses, and a lower induction upon wounding than wax deficient mutant plants, suggesting a trade-off between the production of wax and JA-mediated defenses.

Both cuticular waxes and JA are derived from fatty acids (Kachroo and Kachroo, 2009). On one hand, cuticular wax is a mixture of very-long-chain fatty acid derivatives formed upon the elongation of C_16_ and C_18_ acyl-CoAs (Samuels et al., 2008; Lewandowska et al., 2020). On the other hand, JAs are oxylipins derived from enzymatic or autoxidation of fatty acids (Borrego and Kolomiets, 2016). The *gl8/Ye478* mutant had higher amounts of JA biosynthesis precursors, JAs, and downstream metabolites. This effect was further found to be associated with higher expression of marker genes involved in the JA-biosynthesis pathways gene expression levels and increased concentrations of proteinase inhibitors (Liu et al., 2021), flavonoids (Treutter, 2006), and the benzoxazinoid HDMBOA-Glc (Glauser et al., 2011). The application of a JA biosynthesis inhibitor on mutant plants decreased JA-related defense marker genes in *gl8/Ye478* mutant plants and therewith increased the performance of the insect. Interestingly, this effect was not observed in WT plants, possibly due to the different redox states between *gl8* mutant and WT plants based on our transcriptome analysis. Besides inhibiting JA biosynthesis (Andrade et al., 2017; Zhang et al., 2020), SHAM also has functions of inhibiting the cytochrome pathway and stimulating quinol oxidizing alternative oxidase (AOX). in maize plants (Kolarovic et al., 2006). Altogether, our data demonstrate a trade-off between wax production and JA-mediated defenses, which in turn affect the performance of the herbivore.

Our study shows that *ZmGL8* modulates lipid metabolism and is a key regulator of the balance between wax and JA-derivative synthesis in maize plants. Linoleic and linolenic acids are important precursors for the JA-biosynthesis pathway, and galactolipids MGDG and DGDG are major first precursors for JA biosynthesis (Yu et al., 2020). Galactose liposynthases MGD and DGD are key enzymes for MGDG and DGDG accumulations. The *dgd1* Arabidopsis mutant has much lower concentration of DGDG, but the lipoxygenase-related gene expression levels and OPDA, JA and JA-Ile concentrations are significantly up-regulated (Kelly et al., 2016; Lin et al., 2016). These studies indicate the down-regulation of *ZmDGD1* probably can induce JA accumulation in *gl8/Ye478* mutant, which was consistent with our findings. However, we did not find a clear difference in MGDG and DGDG accumulation between *gl8/Ye478* mutant and WT plants, which is probably because of the fast transformation and the complex structures of galactolipids. Additionally, we suppose that lysophospholipids, PA and FFAs, which accumulated more in *gl8/Ye478* mutant plants, may serve as second messengers to promote herbivore chemical resistance. In our study., we found a clear increase of lysophospholipids and PA with C_18_ structure in *gl8/Ye478* mutant, which are known as key lipid signaling molecules. Previous studies showed that lysophospholipids can activate PLD and many other resistance-related enzymes (Okazaki and Saito, 2014), and PA and oleic acids can also stimulate the JA biosynthesis and signaling pathways (Scherer, 2010; Yu et al., 2010; Mandal et al., 2012). Our results are in line with previous evidence that indicates a deficiency of wax biosynthesis can improve JA concentrations by remodeling plant lipid metabolism. Wan *et al*. found that cuticular wax deficiency improved the C_36_ structure lipids and gene expression levels of related hydrolases in orange peels, which is speculated as the reason for improvement of JA accumulation (Wan et al., 2020). *GhKCS13* encodes VLCFAs biosynthesis enzyme in the ER and epidermal cells, with a potential function for regulating cotton cold resistance by changing lipid components, lipid signaling molecules and JA amounts (Wang et al., 2020).

The trade-off between wax synthesis and JA-related defenses was conserved between maize and Arabidopsis. The maize *gl8* and Arabidopsis *kcs1* mutant plants show similar patterns for wax deficiency mediated herbivore responses, which is probably because both of them function in fatty acid elongation cycles from C_16_/C_18_ acyl-CoAs (Lewandowska et al., 2020). Acyl-CoA elongase is a complex of several enzymes, including ketoacyl-CoA synthases, such as FAE1, KCS1and CUT1 in Arabidopsis, which can generate *β*-ketoacyl-CoA, while *ZmGL8* encodes the *β*-ketoacyl-CoA reductase, which catalyzes the reduction of a ketone group to a hydroxyl group (Xu et al., 2002). Besides, both *ZmGL6* and *ZmGL14* encode proteins for wax transporting and have a slight influence on the biosynthesis of VLCFAs, alkanes or alcohols (Li et al., 2019; Zheng et al., 2019). This probably causes that the *gl6/B73* and *gl14/B73* mutants still have few wax crystals on their leaf surface (Supplementary Figure S9) and have smaller effects than *ZmGL1* and other upstream genes on chemical defense to herbivores. This hypothesis may explain the chemical defense difference among Arabidopsis mutants. In general, Arabidopsis *cer8* mutant showed lower defense responses to herbivore-induction, which is probably because of *CER8* function on VLCFA biosynthesis and transporting to plasma membrane (Lu et al., 2009) (Figure 5). Together, wax biosynthesis gene functions play a vital role in the conservation of trade-off between wax synthesis and JA-related defenses of C16:3 and C18:3 plants, even they have different JA biosynthesis pathways. A better understanding of the molecular and genetic basis of chemical responses manipulated by wax biosynthesis will be required to disentangle these hypotheses.

To minimize defense costs, plants have evolved different strategies to modulate their complex defense networks against herbivores, such as the trade-off between physical and chemical defenses (Cornelissen et al., 2009; Read et al., 2009; Descombes et al., 2019). Wax characters and their close relationship to JA biosynthesis are probably resulted from the long history of plant-herbivore coevolution (Shepherd and Griffiths, 2006). The scenario may be that as the cuticular wax can prevent insect feeding, plants tend to save the energy on chemical defenses. However, the insects have been gradually increasing their resistance to waxes, which could be a risk for wax-producing plants. As many trade-offs between physical and chemical defenses have been reported, such as phenolic concentrations and fiber height (Cornelissen et al., 2009; Moles et al., 2013), we speculate an overlooked but widespread plant-herbivore arms-races that driven by not only one but several plant defense traits. The illustration of multiple plant traits driven plant-herbivore arms-races probably can write a new chapter in plant-herbivore coevolution, and supply an innovative step forward for agronomic trait selection in sustainable agriculture.

In conclusion, our study reveals the pivotal role of maize GL8 and Arabidopsis KCS1 in mediating trade-offs between cuticular wax biosynthesis and JA-mediated chemical defenses to herbivores. The environmental and genetic mechanisms regulating the plant’s decision to invest its resources in wax production or JA-defenses remain to be investigated. Yet, our finding contributes to a better understanding of plant-herbivore coevolution and opens new avenues for herbivore management.

## Methods

### Plant and Insect Resources

Maize (*Zea mays* L.) genotypes Ye478, B73, *gl1/B73*, *gl6/B73*, *gl8/B73*, *gl14/B73* and 2,000 materials of Ye478 ethyl methane sulfonate (EMS) mutagenized mutant library were used in this study. The EMS mutant library was generated by Fanjun Chen from China Agricultural University. Maize seeds were sowed in pots (7.5 × 6.0 × 7.5 cm) with soil and roseate (3:1, v/v) and grown in a greenhouse (28 ± 3°C day, 22 ± 3°C night, 50 - 60% relative humidity, 14/10 h light/dark) for 14 days before experiments.

*Arabidopsis thaliana* genotypes *cer1*, *cer3*, *cer8*, *kcs1* and Col-0 were bought from AraShare service center (https://www.arashare.cn/index/Product/index.html). Arabidopsis seeds were planted on ½MS medium (pH 5.8; Sigma-Aldrich) and were stratified at 4 °C for 3 days in the dark, and then transferred to controlled growth chambers (22 °C and 100 μmol m^-2^ s^-1^ light intensity under 16/8 h light/dark conditions). Seven days later, seedlings were transferred to soil in pots (7.5 × 6.0 × 7.5 cm). Arabidopsis seedlings were used 21 days after sowing.

Fall armyworm (*Spodoptera frugiperda*) individuals were collected in Zhengzhou, Henan province of China (34°25’N,113°43’E) in 2020, and reared on an artificial diet as described before (Maag et al., 2014). Fall armyworm larvae were starved for 10 hours prior use in experiments. Oral secretion (OS) of *S. frugiperda* was collected from third-instar larvae that were fed on maize leaves and diluted 1:1 with sterilized Milli-Q water (Millipore) before use. To mimic herbivore feeding, the lower epidermis of third fully unfolded maize leaf was damaged on both sides of the central vein by knife blades until the mesophyll tissue was removed. The damaged area was around 3-4 cm^2^ and then 10 μL OS was added on the damaged parts.

### Cuticular Wax Phenotyping

The phenotyping of cuticular wax deficient genotypes in maize and Arabidopsis was conducted by Field Emission Scanning Electron Microscopy (FE-SEM). Briefly, some water was sprayed on the plants. Plants that displayed a water-holding phenotype were further used for SEM analysis. The second leaf of maize plants and the youngest *A. thaliana* leaf was cut into 1×1 cm small pieces and then characterized with a FE-SEM (Quanta 250, FEI, USA).

### Herbivore Bioassays

Herbivore damage was tested *in vitro*. Briefly, one larva was placed into a Petri dish (90 mm diameter) containing a plant leaf (n=14-15). Third maize leaves or youngest *A. thaliana* leaves were used in the assay. After 8 hours, all leaves were collected to measure the damage area with ImageJ.

Herbivore performance was assessed *in vivo*. Four plants (all maize or all Arabidopsis) were grown per pot (n=20). Three third-instar pre-weighed larvae were placed on all plants. Plants in each pot were covered by an oven bag. After 4 days, larvae were collected and weighed again to calculate the growth rate.

### Phytohormone Quantification

Non-induced and OS-induced maize and *A. thaliana* leaves were collected 1.5 h after induction and ground in liquid nitrogen (n=4). OPDA, JA, JA-Ile, SA, ACC and ABA were extracted with mixed-mode strong cation exchange-mixed-mode weak anion exchange (MCX-WAX) solid-phase extraction (SPE) progress as described by Xin et al. (Xin et al., 2020). Extracted solvents with isotopically labeled standards inside (100 ng for d_5_-JA, d_6_-ABA, d_2_-IAA, d_4_-SA, d_5_-OPDA, d_4_-ACC and ^13^C_6_-JA-Ile) were analyzed by ultrahigh-performance liquid chromatography-tandem mass spectrometry (UPLC-MS/MS) following the previous described procedures (Glauser et al., 2014).

### Flavonoid and Benzoxazinoid Analyses

Flavonoid contents in OS-induced and non-induced maize leaves were analyzed by spectrophotometry. The third leaf of control or treated maize plants was collected 0 h or 1.5 h after induction, and five leaves from five plants were pooled as one sample (n=3 replicates of five plants each). Then, the samples were freeze dried by vacuum freeze drier (Scanvac coolsafe 110-4, Labogene, Denmark) immediately and grounded into powder in liquid nitrogen. Aliquots of 100 mg leaf powder were extracted by 70% ethanol and then fully mixed under 30 Hz ultrasonic waves for 1 h at 40°C. The samples were then filtered through a filter paper and measured by ultraviolet spectrophotometer at 495 nm. A rutin standard curve was used for quantification, as described by (Chen et al., 2016).

Benzoxazinoids were measured 1.5 h after OS induction in the third leaf of induced and non-induced maize plants. Pools of three leaves (from three plants) were used (n=4 of 3 plants each). All samples were ground in liquid nitrogen. Benzoxazinoids were extracted by adding 500 μL 50% MeOH + 50% H_2_O + 0.5% formic acid to aliquots of 50 mg leaf powder. The extracts were vortexed for 1 min and centrifuged twice at 17,000 g, at 4 °C. The supernatants were then analyzed by UPLC-MS/MS system as described before (Zhang et al., 2019).

### Gene Expression Analysis

Expression levels of herbivore resistance related genes in maize and *A. thaliana* were assessed by quantitative real-time polymerase chain reaction (qRT-PCR) (Hu et al., 2019). Leaf samples were collected 0 and 1.5 h after OS induction. All samples were ground in liquid nitrogen. Total RNA was isolated by TRIzol Reagent (Life Technologies, USA) following the manufacturer’s protocol. The obtained RNA was reverse transcribed with Revert Aid First Strand cDNA Synthesis Kit (Vazyme, Nangjing, China) according to the manufacturer’s instructions. qRT-PCR analyses were conducted on a LightCycler® 96 System (Roche Diagnostics GmbH, Mannheim, Germany) using the SYBR Green I Master Mix (Vazyme, Nanjing, China). Ubiquitin genes of maize *ZmUbiquitin2* and *AtUBQ10* were used as internal standards to normalize cDNA concentrations, and the relative gene expression levels were calculated based on 2^−ΔΔCt^ method (Wang et al., 2019). All primers used for qRT-PCR are listed in Supplemental Table S2.

### Transcriptome Analysis

RNA-seq analysis was performed on OS-induced and non-induced Ye478 and *gl8/Ye478* maize plants. Five leaf samples were pooled together (n=3 pools of 5 plants). HiSeq 2500-PE125 platform (Illumina) at a 6G depth was conducted. The raw data was filtered by Cutadapt to remove reads with 3’ adaptors and the low quality. Then, the clean reads were mapped to the maize B73 reference genome (RefGen_V4) with HISAT2 (Siren et al., 2014). The Read Count of each gene was calculated by HTSeq and then estimate abundance with STRINGTIE (Pertea et al., 2016). The differentially expressed genes (DEGs) was identified by DESeq (Love et al., 2014), Blast2go and KAAS were used for Kyoto Encyclopedia of Genes and Genomes (KEGG) analysis. The heat maps for herbivore resistance genes were performed by R package ‘pheatmap’.

### Lipidomic Analysis

Plant lipid profiles were characterized in pools of six individual samples (n=5 of third leaves six plants each). Aliquots of 20 mg were used for extraction by adding 1 mL extraction buffer (methyl tert-butyl ether: MeOH = 3:1, V/V), vortexing for 30 min, adding 300 μL sterilized Milli-Q water, and vortexing again for 1 min. The extracts were then kept at 4°C for 10 min before centrifugation at 12000 r/min, 4°C for 3 min. Four hundred microliter of the supernatant was transferred into 1.5 mL eppendorf tubes and dried at 20°C for 2 h. The pellet was resuspended in 200 μL lipid resolving solution (acetonitrile: isopropanol = 1:1, V/V) and centrifuged again at 12000 r/min, 4°C for 10 min. The resulting supernatant was used to analyzed lipids by UPLC (ExionLC™ AD, SCIEX, USA) connected with tandem mass spectrometry (MS/MS) (QTRAP®6500^+^, SCIEX, USA). Briefly, a Thermo Accucore™C30 column (2.6 μm, 2.1 mm×100 mm i.d, 45°C) and binary solvent system was used. Solvent A was 40% acetonitrile and solvent B was acetonitrile/isopropanol (1:9, V/V), and both solvents contained 0.1% formic acid and 10 mmol/L ammonium formate. A linear gradient profile with solvent A and B (flow rate = 0.35 mL/min.) was applied: 0-2 min, 80% A; 2-4 min, 70% A; 4-9 min, 40% A; 9-14 min, 15% A; 14-15.5 min 10% A; 15.5-17.5 min, 5% A; 17.5-20 min, 80% The injection volume was of 2 μL. MS conditions were as follows: electrospray ionization (ESI) temperature 500 °C; MS voltage at positive mode 5.5 kV, and negative mode −4.5 kV; gas 1 and 2 (nitrogen), 50 psi; and curtain gas (nitrogen), 35 psi. The scan range of *m/z* was based on optimized decluttering potential and collision energy. The lipid qualitative analysis was based on Metware Database and the quantitative analysis was conducted by multiple reaction monitoring mode in MS system (Lu et al., 2019).

### Inhibitor Treatments

The suppression of JA biosynthesis was obtained by using the lipoxygenase inhibitor salicylhydroxamic acid (SHAM) in *gl8/Ye478* mutant and WT maize plants. For each plant, 5 mL of 400 mΜ SHAM in MilliQ water was supplied around the root, and 5 mL sterilized Milli-Q water was supplied to control plants. The herbivore feeding and performance bioassays were conducted after 24 h of SHAM and water supplement.

### Statistical Analysis

Statistical analyses were conducted in R 4.0.5. The normality of the residue distribution and the heteroscedasticity of the data were verified prior analyses. As the assumptions for parametric tests were fulfilled, herbivore damage and performance, gene expression levels, secondary metabolite concentrations were analyzed by Student’s *t*-tests or Analyses of Variance (ANOVA). For pairwise comparisons, Least Squares Means (LSMeans) and FDR-corrected post hoc tests were conducted. The R packages involved in all statistical analyses were “car”, “lsmeans”, and “RVAideMemoire” (Benjamini and Hochberg, 1995; Lenth, 2016; Herve, 2022).

## Acknowledgements

We thank Prof. Yuan Zheng (Henan University) for his valuable feedback, Lingyu Mi for the technical assistance in electron microscopy, Prof. Jun Zheng (Institute of Crop Science, Chinese Academy of Agricultural Sciences) for providing *gl6* mutant seeds, and Fanjun Chen for supplying the EMS mutant library. This work was supported by the National Science Foundation of China to Chun-Peng Song (U21A20206), the National Natural Science Foundation of China (32102187) to Xi Zhang, the Project of Sanya Yazhou Bay Science and Technology City (SCKJ-JYRC-2022-78) to BaoZhu Li and the Program of Introducing Talents of Discipline to Universities (111 Project, number D16014) to Shutang Zhou and Xi Zhang.

## Author contributions

Chun-Peng Song, Xi Zhang and Shutang Zhou conceived and designed the research. Xi Zhang and Jiong Liu wrote the original draft. Chun-Peng Song, Shutang Zhou, Xi Zhang, Christelle AM Robert and Jiong Liu revised the manuscript. Xi Zhang, Jiong Liu, Lu Li performed all experiments with the help of Zhilong Xiong, Wenjie Chen, Jiasheng Bi, Guanqing Zhai, Shan He, Hui Zhang. Baozhu Li, Siyi Guo and Jieping Li. Jieping Li and Shutang Zhou supported the production of mutant lines, the map-based cloning work and insect resources. Xi Zhang, Jiong Liu, Lu Li and Baozhu Li analyzed the data.

---

**Supplemental Figure S1.**
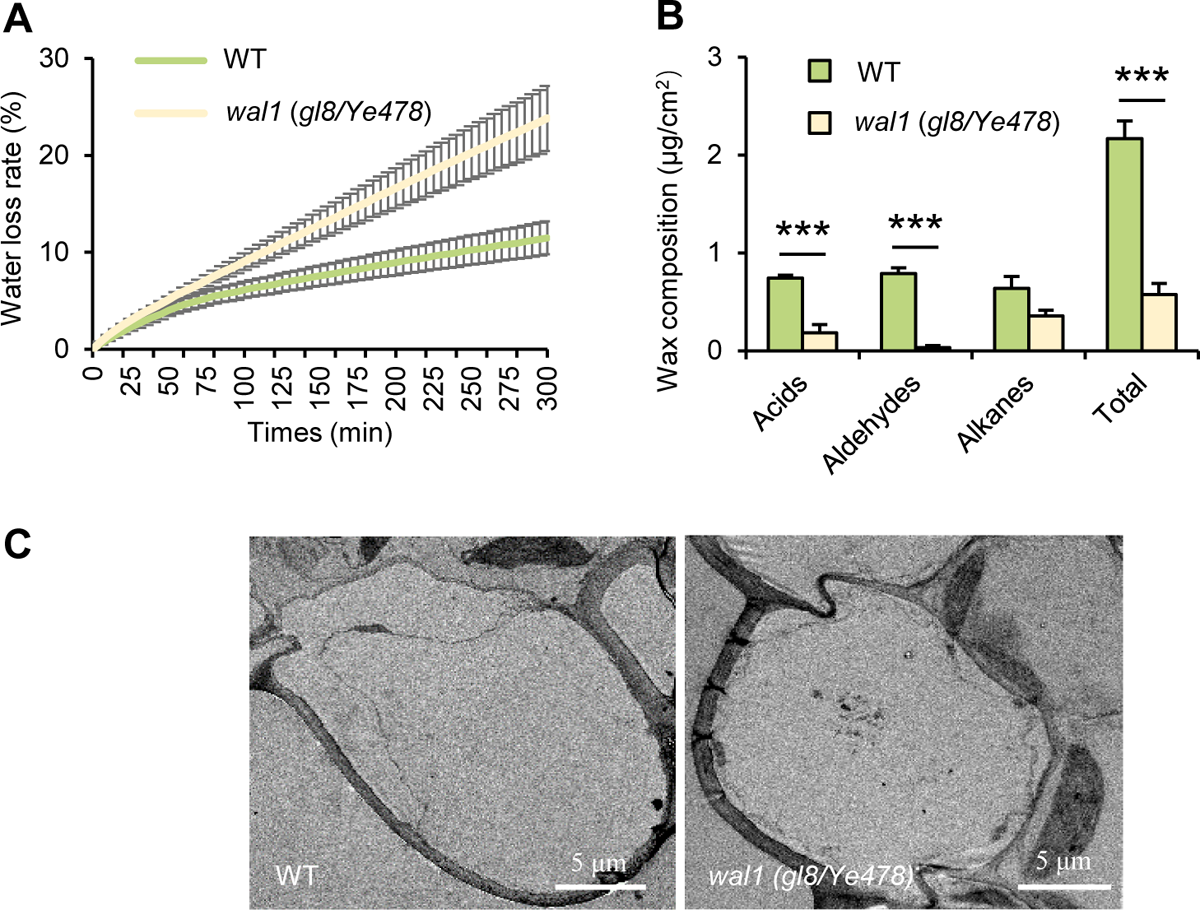
Phenotyping of *wal1* (*gl8/Ye478*) mutant. **(A)** The water-loss rate of WT and *gl8/Ye478* mutant. **(B)** Total leaf cuticular wax load and wax composition of WT and *wal1* (*gl8/Ye478*) mutant (Mean ± SE, n = 4 biological replicates, with eight plants pooled per replicate). Asterisks indicate statistically significant differences between *wal1* (*gl8/Ye478*) mutant and WT maize plants (Student’s *t*-test, *\*P* < 0.05, *\*\*P* < 0.01, *\*\*\*P* < 0.001). **(C)** Transmission scanning electron microscopy analysis of cross-sectional microstructure of epidermal cells in WT and *wal1* (*gl8/Ye478*) mutant. Bar = 5 μm.

**Supplemental Figure S2.**
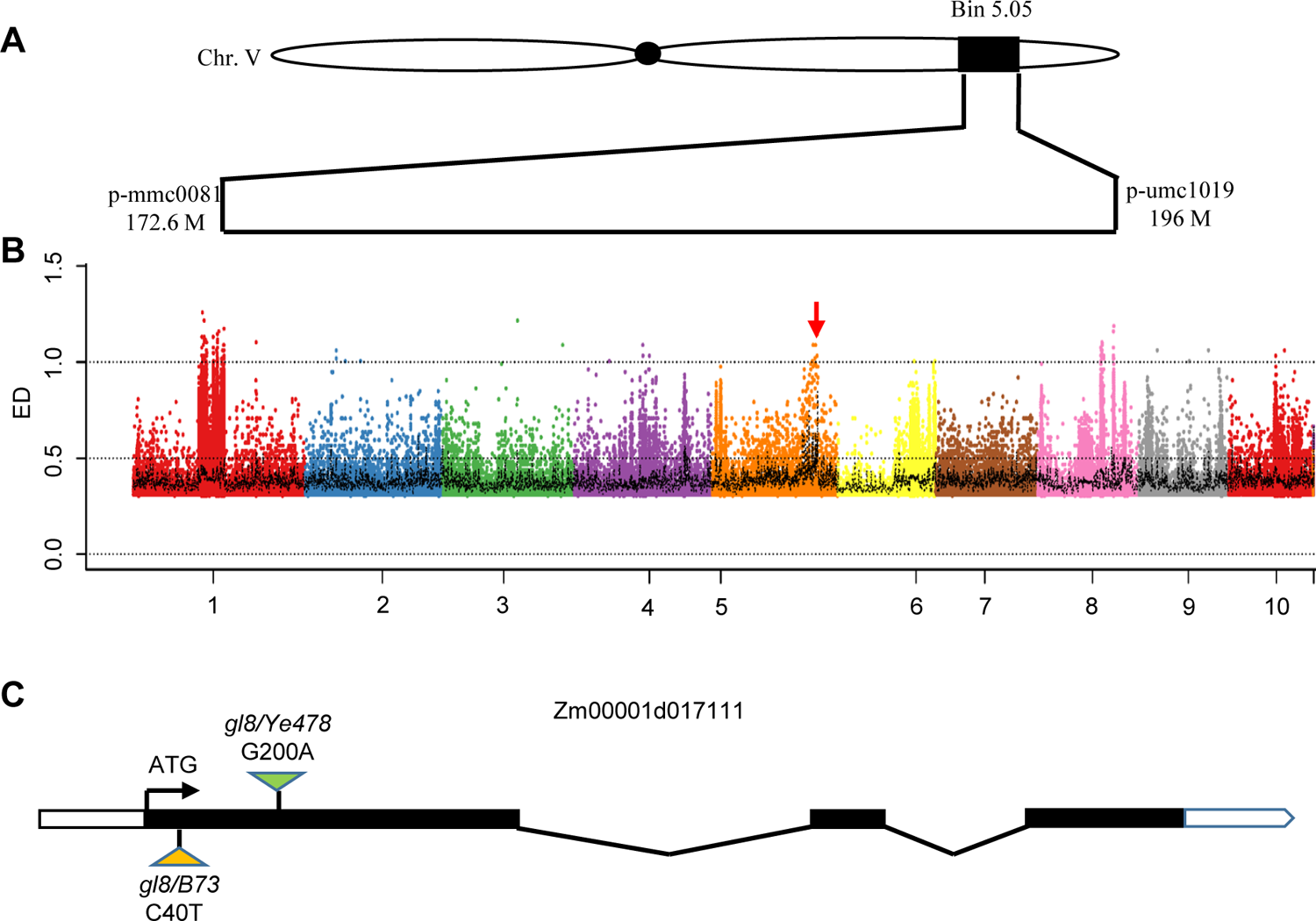
Map-based cloning and identification of *ZmGL8.* **(A)** *GL8* was mapped to bin 5.05 of chromosome V, between the p-mmc0081 and p-umc1019 SSR markers, a 24.6 Mb region. **(B)** Through the whole genome sequencing Euclidean distance analysis, it can be determined that the target gene causing non-synonymous mutation in the range of 172.6 M-196 M at the end of chromosome V is *ZmGL8* (Zm00001d017111). **(C)** The mutation site of *gl8/Ye478* was mutated from G to A at the 200^th^ base from the start codon, causing a non-synonymous mutation. The mutation site of *gl8/B73* is the mutation of C to T at the 40^th^ base from the start codon, forming a stop codon.

**Supplemental Figure S3.**
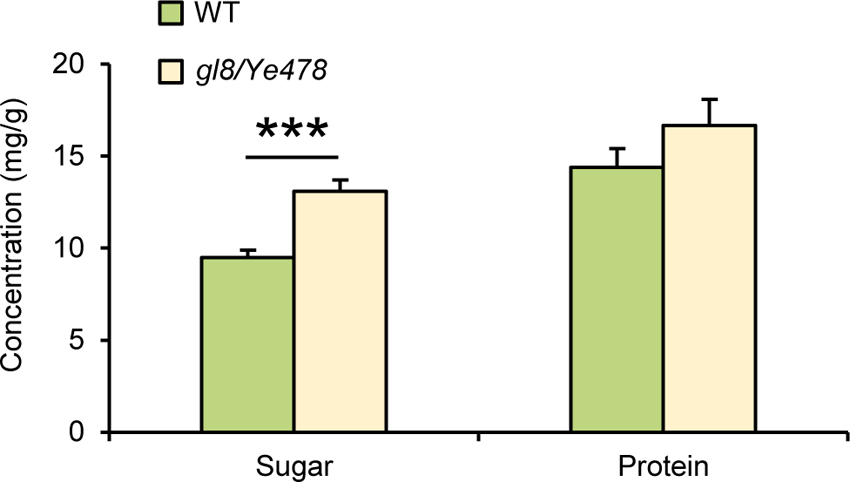
Nutrient levels of *gl8/Ye478* mutant and WT maize plants. Average concentrations of total soluble sugar (Mean ± SE, n = 6) and protein (Mean ± SE, n = 6) of *gl8/Ye478* mutant and WT maize plants. Asterisks indicate statistically significant differences between *gl8/Ye478* mutant and WT maize plants (Student’s *t*-test, *\*\*\*P* < 0.001).

**Supplemental Figure S4.**
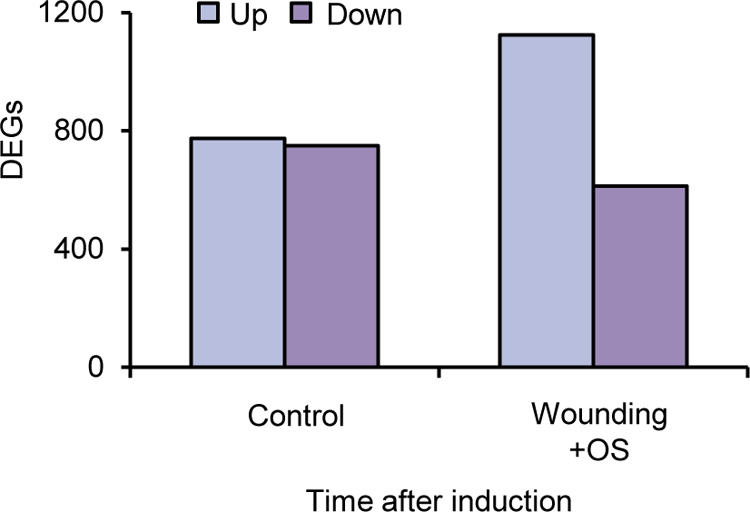
The number of different expression gene numbers of *gl8/Ye478* mutant and WT maize plants by RNA-Seq.

**Supplemental Figure S5.**
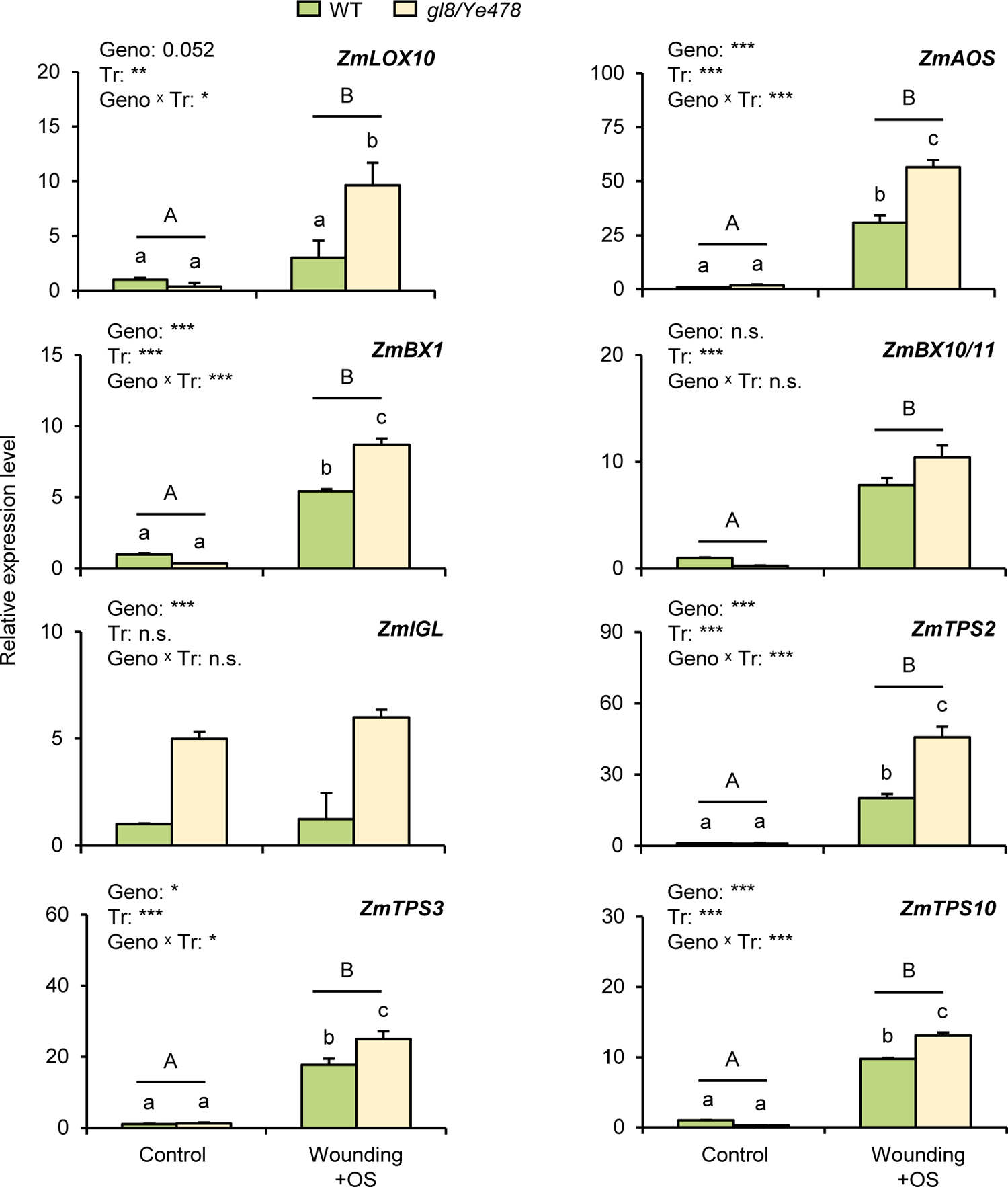
The expression of insect resistance related genes in maize was increased after herbivore induction treatment. Gene expression levels (Mean ± SE, n = 3) of *ZmLOX10*, *ZmAOS*, *ZmBX1*, *ZmBX10/11*, *ZmIGL*, *ZmTPS2*, *ZmTPS3* and *ZmTPS10* of *gl8/Ye478* mutant and WT maize plants under control and wounding + OS induction treatments through qantitative real time polymerase chain reaction (qRT-PCR) analyses. Asterisks indicate significant differences between genotypes, treatments or the two factor interactions (*\*P* < 0.05, *\*\*P* < 0.01, *\*\*\*P* < 0.001). Lower case letters indicate the significant differences of pairwise comparisons, and uppercase letters indicate significant differences between control and wounding + OS treatments.

**Supplemental Figure S6.**
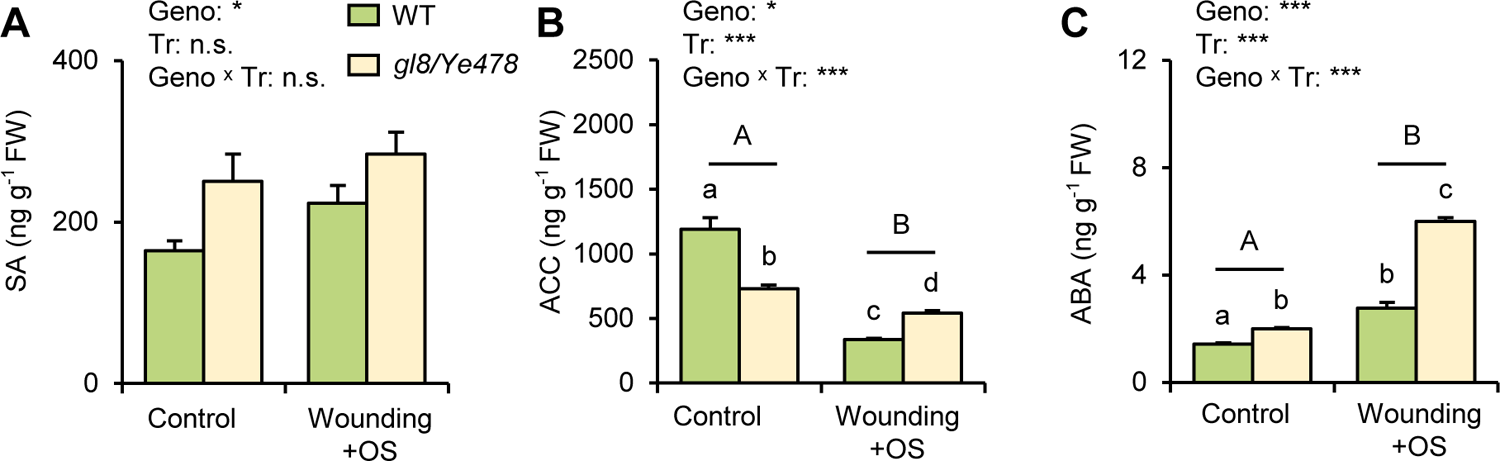
Phytohormone analysis of *gl8/Ye478* mutant and WT maize plants under control and herbivore induction treatments. **(A)** Salicylic acid (SA), **(B)** 1-aminocyclopropane-l-carboxylic acid (ACC, a biosynthetic precursor of ethylene) and **(C)** abscisic acid (ABA) average concentrations (Mean ± SE, n = 4) of *gl8/Ye478* mutant and WT maize plants under control and wounding + OS induction. Asterisks indicate significant differences between genotypes, treatments or the two factor interactions (*\*P* < 0.05, *\*\*\*P* < 0.001). Lower case letters indicate the significant differences of pairwise comparisons, and uppercase letters indicate significant differences between control and wounding + OS treatments.

**Supplemental Figure S7.**
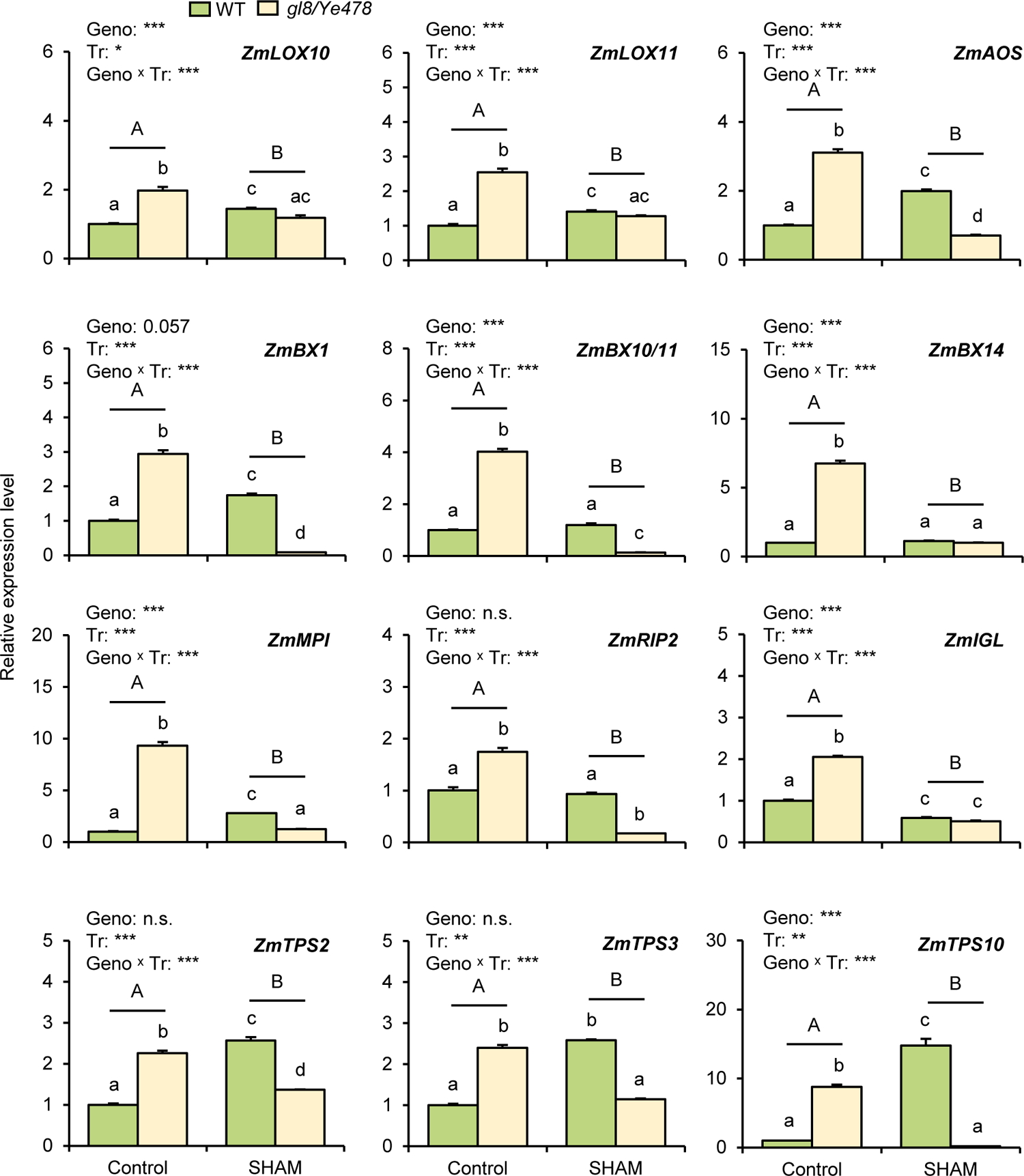
JA biosynthesis inhibitor SHAM treatment inhibited the expression of maize insect resistance related genes in *gl8/Ye478* mutant. Gene expression levels (Mean ± SE, n = 3) of *ZmLOX10*, *ZmLOX11*, *ZmAOS*, *ZmBX1*, *ZmBX10/11*, *ZmBX14*, *ZmMPI*, *ZmRIP2*, *ZmIGL*, *ZmTPS2*, *ZmTPS3* and *ZmTPS10* of *gl8/Ye478* mutant and WT maize plants treated with JA biosynthesis inhibitor SHAM through qRT-PCR analyses. Asterisks indicate significant differences between genotypes, treatments or the two factor interactions (*\*P < 0.05*, *\*\*P < 0.01*, *\*\*\*P < 0.001*). Lower case letters indicate the significant differences of pairwise comparisons, and uppercase letters indicate significant differences between control and wounding + OS treatments.

**Supplemental Figure S8.**
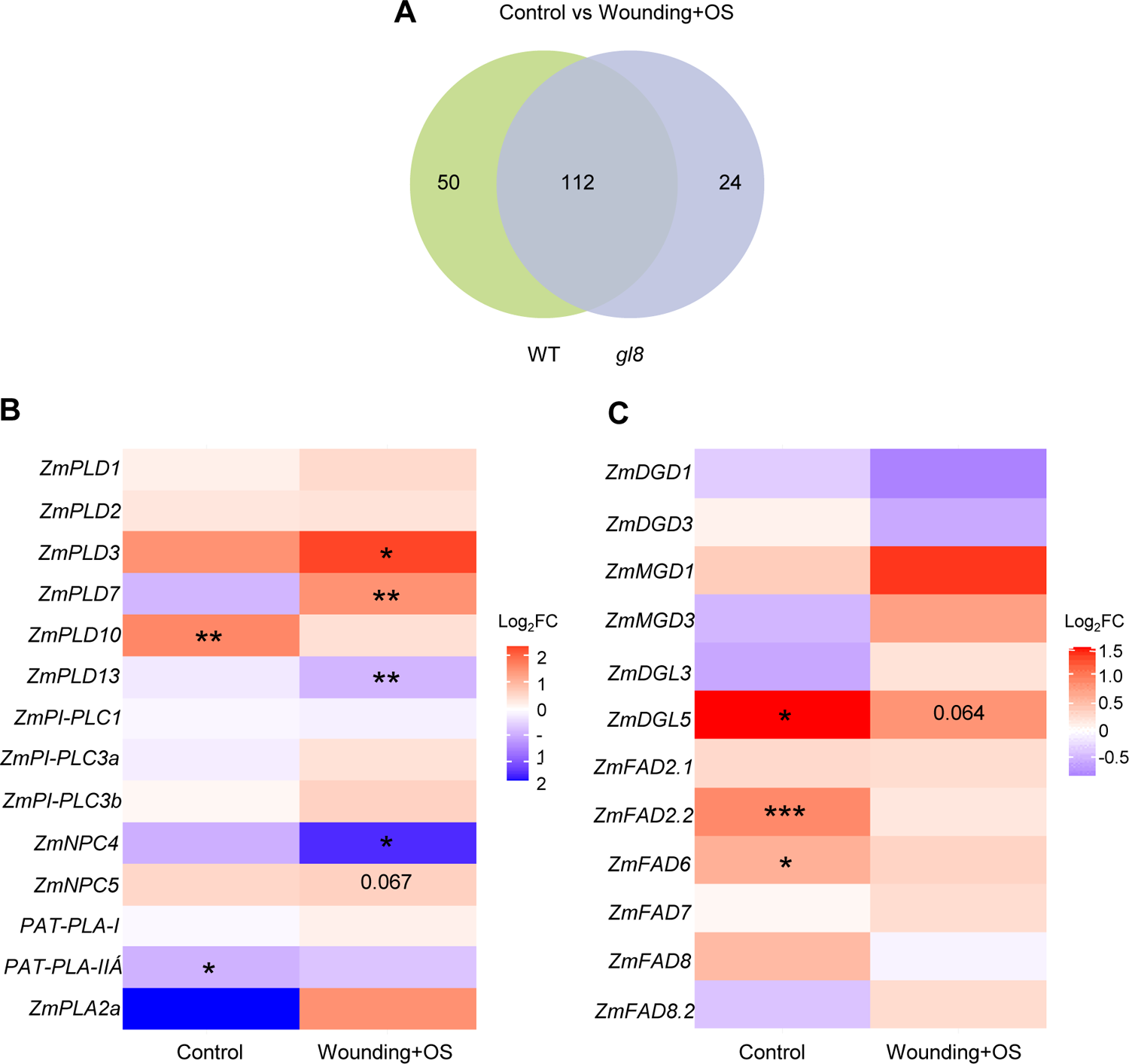
Lipinomic analysis between *gl8/Ye478* mutant and WT maize plants under control and herbivore induction treatments. **(A)** Venn diagram analysis of different expressed gene numbers of *gl8/Ye478* mutant and WT maize plants under control and wounding + OS treatments. **(B)**-**(C)** Heat map of genes related to *GL8* modulated lipid metabolism in *gl8/Ye478* mutant compared with WT plants (n = 5). The color gradient represents the relative sequence abundance. Numbers in the color key indicate log_2_ fold change (FC).

**Supplemental Figure S9.**
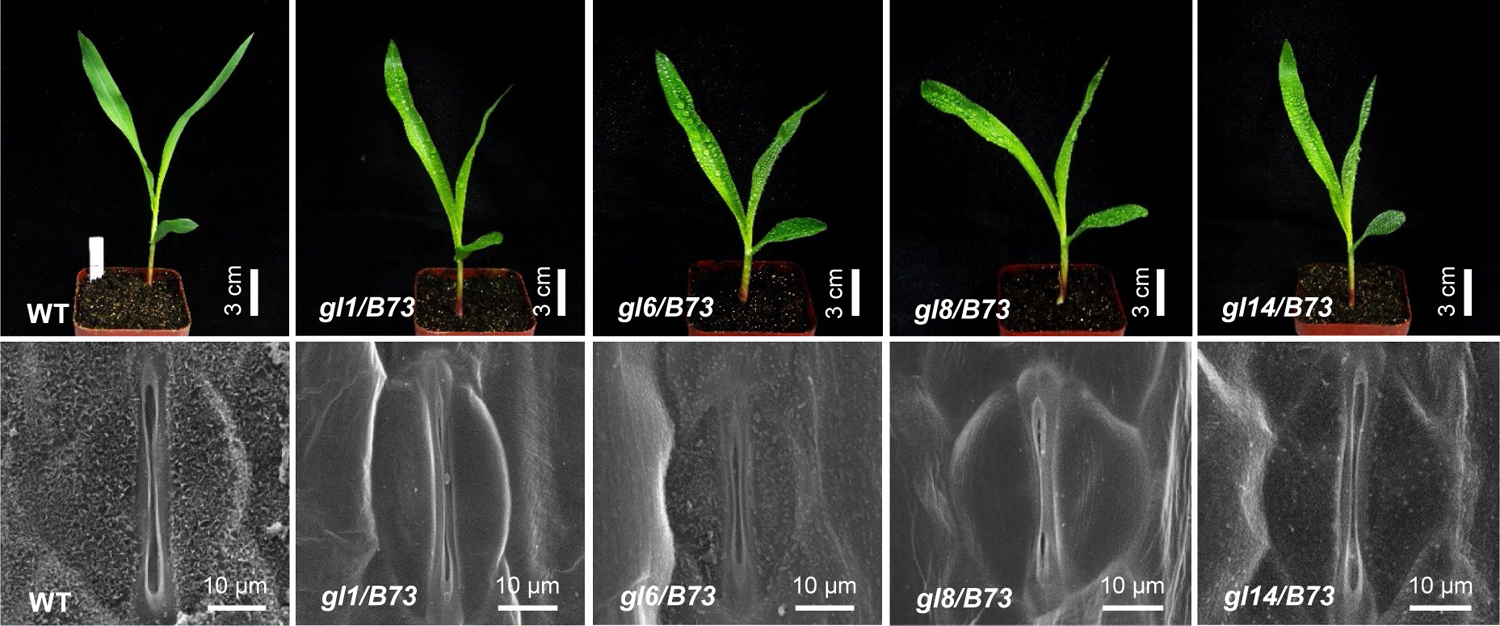
Cuticular wax phenotype of selected maize wax deficiency mutants. **(A)** Phenotype of four *glossy* mutants and WT maize plants in B73 background after water spraying. Bar = 3 cm. **(B)** Adaxial leaf cuticular wax accumulation on WT and four *glossy* mutant seedling leaves detected via SEM (×6000 magnification, bar = 10 μm).

**Supplemental Figure S10.**
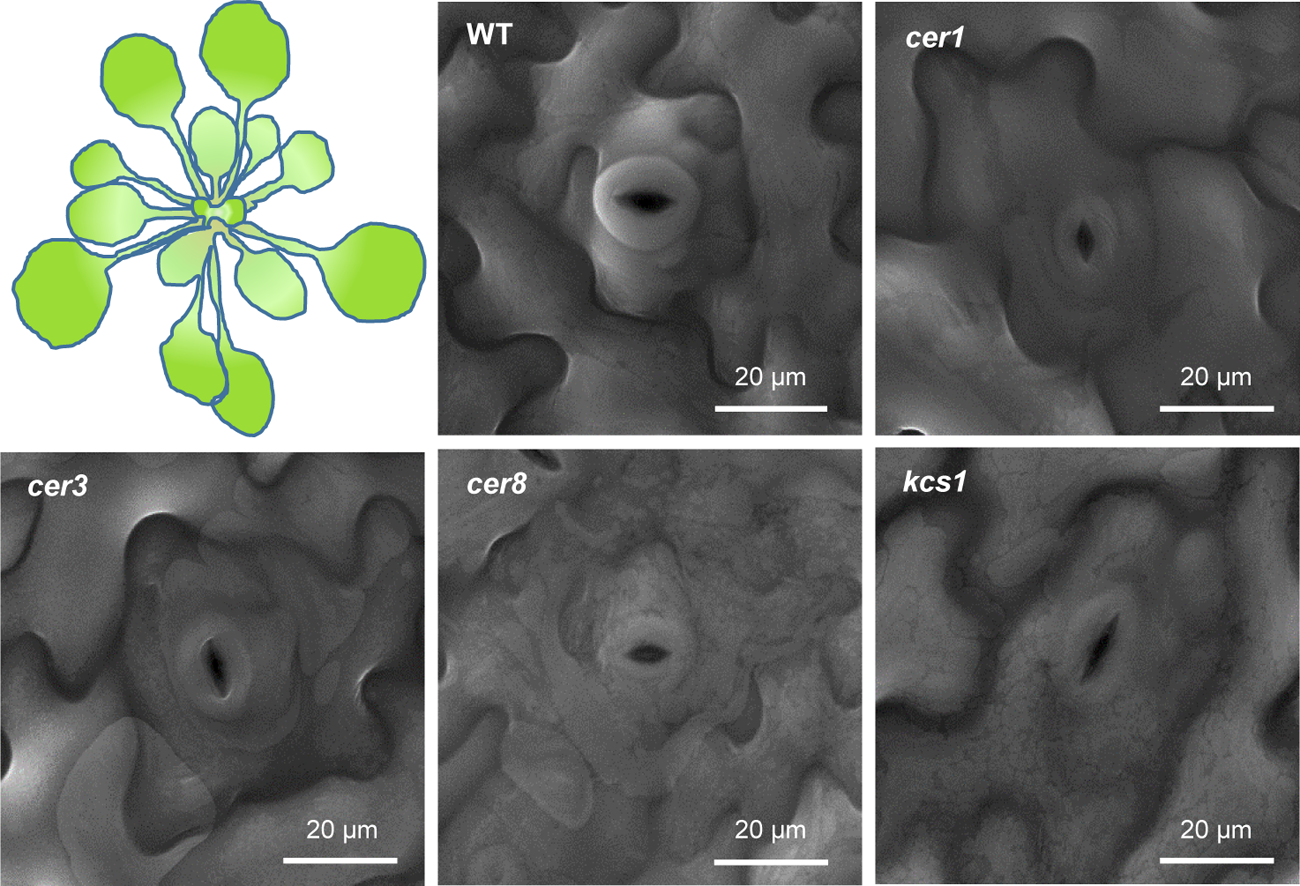
Cuticular wax phenotype of selected *Arabidopsis thaliana* wax deficiency mutants. Adaxial leaf cuticular wax accumulation on WT, *cer1*, *cer3*, *cer8*, and *kcs1* mutants detected via SEM (×4000 magnification, bar = 20 μm).

**Supplemental Figure S11.**
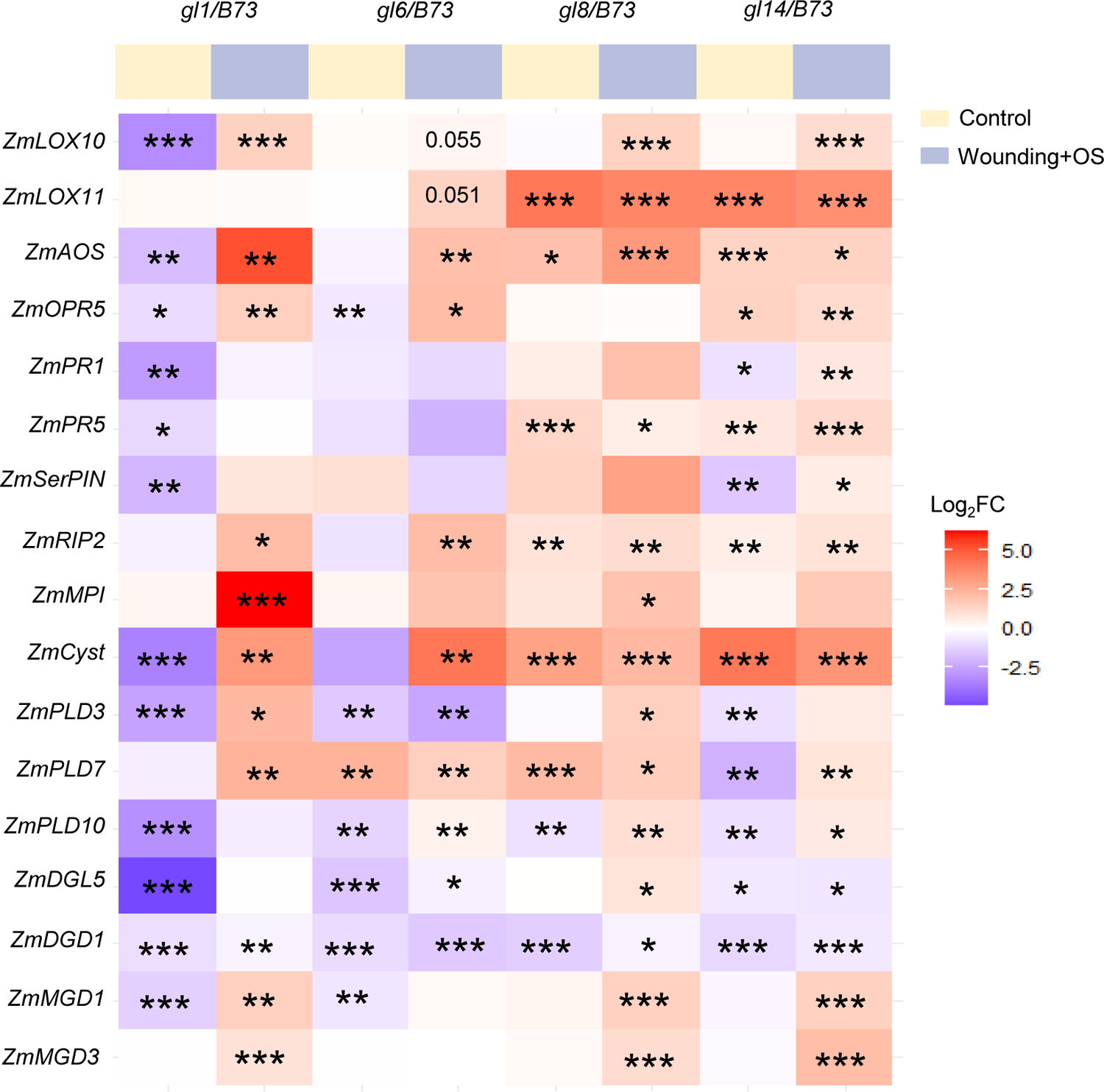
The important insect resistance related genes in maize cuticular wax deficiency mutants were significantly up-regulated compared with wild type. Heat map of genes related to herbivore defenses in wax deficiency mutants compared with WT maize plants in B73 background. The color gradient represents the relative sequence abundance. Numbers in the color key indicate log_2_FC. Asterisks indicate significant differences between genotypes (*\*P < 0.05*, *\*\*P < 0.01*, *\*\*\*P < 0.001*).

**Supplemental Table S1.** The segregation statistics of the F_1_ generation of the cross between *gl8/Ye478* and *gl8/B73*

|  | Normal plant (NP) | Mutant plant (MP) | NP: MP |
| --- | --- | --- | --- |
| Observed | 667 | 203 | 3.29:1 |
| Expected | 653 | 217 | 3:1 |

$\chi^2 = 1.2 < \chi^2_{0.05} = 3.84$ . The difference was extremely significant.

**Supplemental Table S2.**
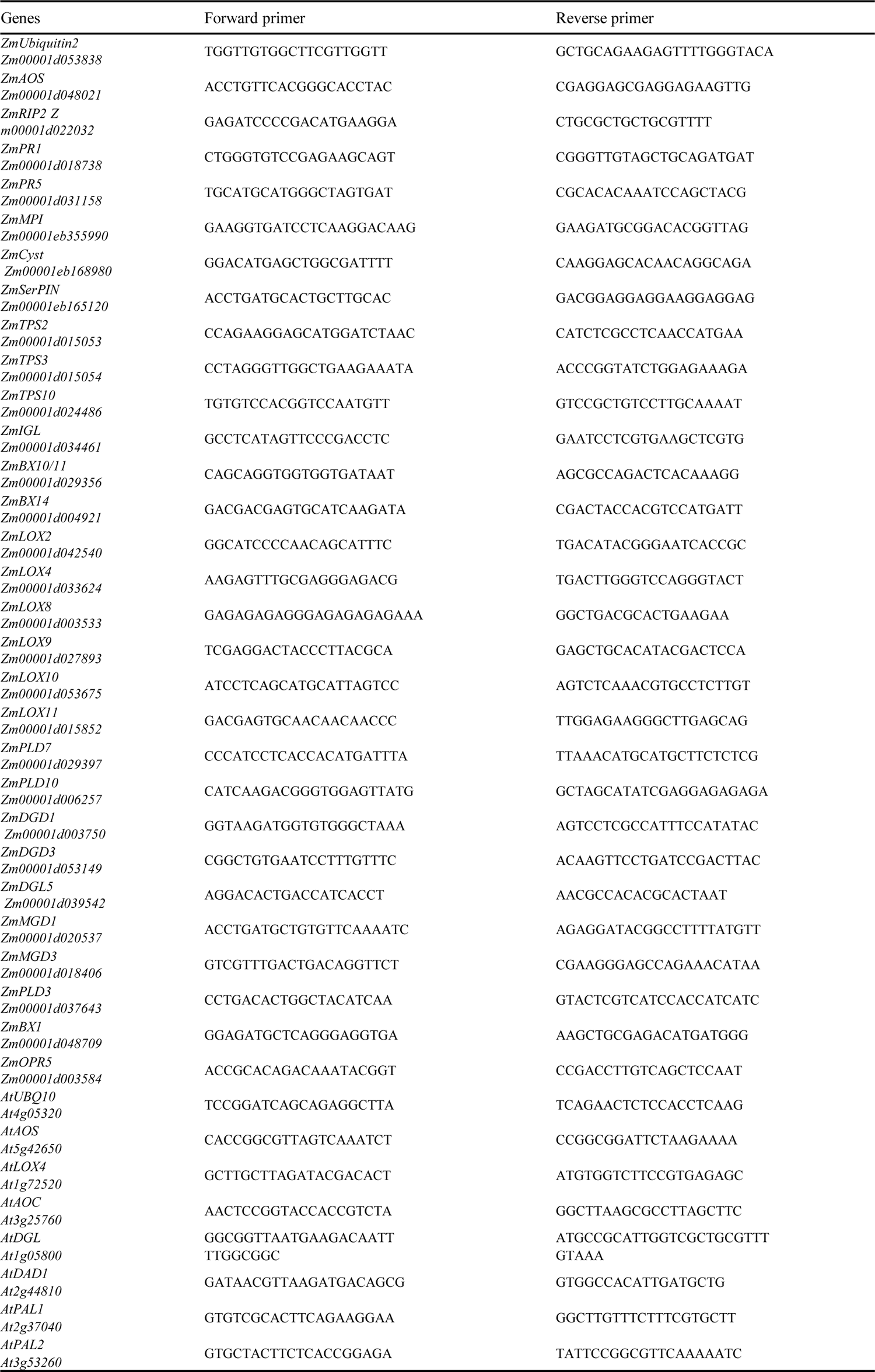
The primers used for qRT-PCR in this study.

**Supplemental Table S3.**
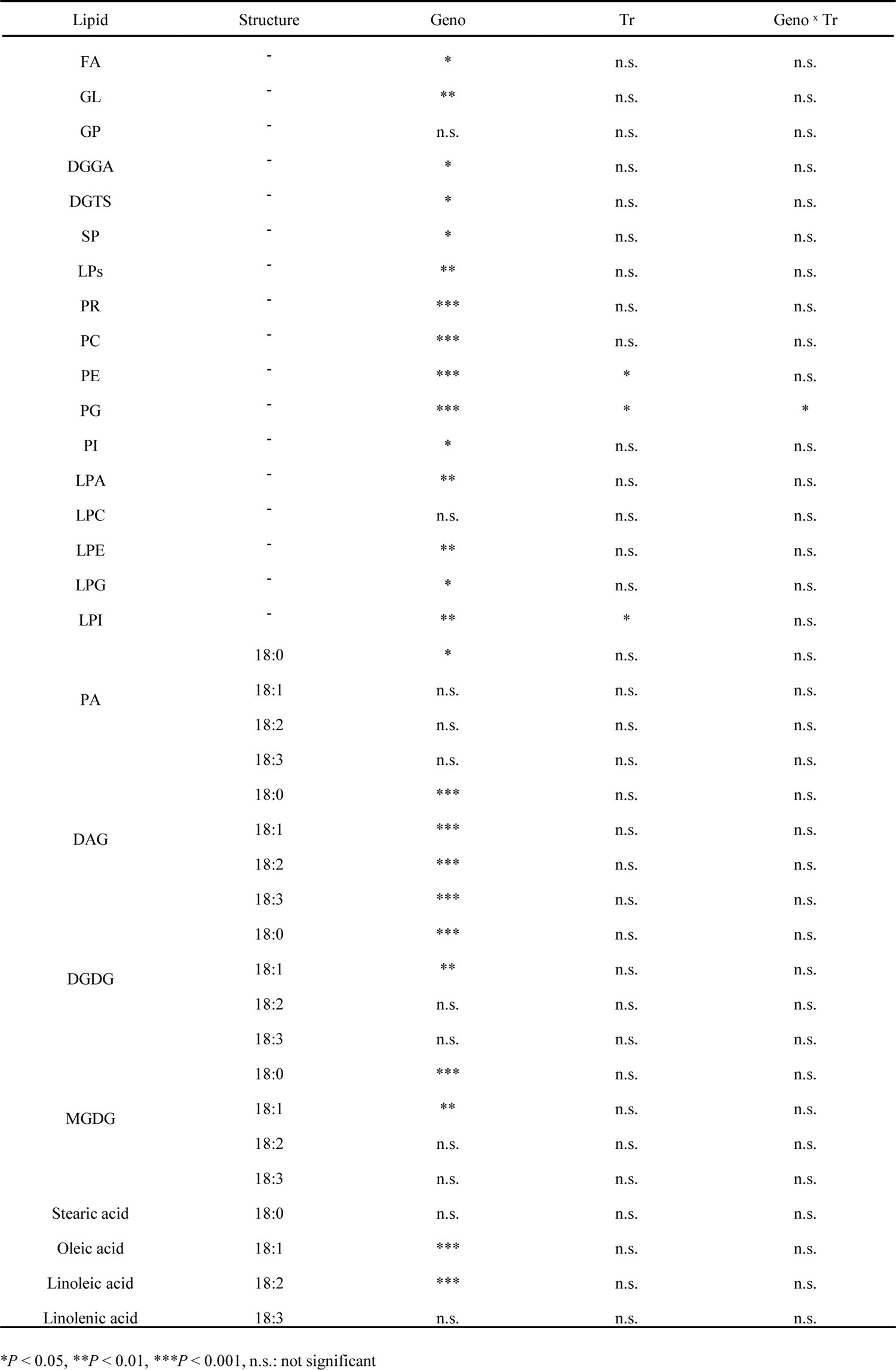
The ANOVA analysis results for lipids shown in Figure 3.

